# VITAL-3D: Volumetric Single-Cell Quantification Reveals Microenvironment-Dependent Drug Responses in Breast Cancer

**DOI:** 10.64898/2026.08.31.748448

**Authors:** Chongguang Jin, Tiffany Chu, Qiming Zhang, Denis Wirtz, Pei-Hsun Wu

## Abstract

Preclinical drug evaluation relies heavily on two-dimensional (2D) monolayer assays, which fail to recapitulate the structural and functional complexity of the tumor microenvironment and may therefore misrepresent therapeutic efficacy. Here, we present VITAL (Volumetric Imaging-based Toxicity and Live Analysis), a high-throughput imaging platform that enables direct single-cell quantification of proliferation and cell death in both 2D and three-dimensional (3D) extracellular matrix (ECM) cultures using a 96-well format. By combining volumetric imaging with automated single-cell analysis, VITAL enables dynamic assessment of drug responses beyond conventional viability assays and EC₅₀ measurements. Using breast cancer cell lines treated with anticancer agents, we systematically compared drug responses between 2D and 3D microenvironments. Although EC₅₀ values were often comparable between culture formats, growth kinetics and concentrations required to induce complete growth arrest or net cell loss differed substantially in 3D cultures. In particular, drug concentrations required to induce net cell loss were consistently higher in 3D, revealing microenvironment-dependent survival responses that were not captured by EC₅₀ alone. Furthermore, clinically expected subtype-specific responses, including tamoxifen sensitivity in ER-positive cells and olaparib sensitivity in BRCA1-mutant cells, were more accurately resolved under 3D culture conditions and extended treatment durations. Together, these findings demonstrate that growth-based, single-cell quantification provides a more comprehensive assessment of therapeutic efficacy than conventional endpoint measurements and establish VITAL as a scalable platform for physiologically relevant preclinical drug screening.

## Introduction

Cancer represents a major global health challenge and is a leading cause of morbidity and mortality worldwide^1, 2^. With the global cancer burden continuing to rise, healthcare systems face increasing pressure to improve cancer prevention, diagnosis, and treatment^1, 3^. There is a continued need to identify more effective anticancer agents^4^. In addition, more than 50% of drugs entering phase III clinical trials fail because of limited therapeutic benefit^5, 6^. Therefore, it is critical to assess drug efficacy using biologically relevant models that more accurately recapitulate the physiological and pathological conditions observed in patients. In vitro models are essential for identifying new anticancer drug candidates and evaluating their efficacy.

*In vitro* models are essential for identifying new anticancer drug candidate and evaluate their efficacy. Conventional drug screening has relied predominantly on two-dimensional (2D) monolayer cultures because of their simplicity, low cost, and compatibility with high-throughput screening^7, 8^. Accumulating evidence demonstrates that the molecular, transcriptional, and functional states of cancer cells differ substantially between 2D and 3D culture systems^9–13^. For example, colorectal adenocarcinoma cells exhibit widespread changes in gene expression when cultured in 3D, while breast cancer cells display alteration in mechanotransduction signaling, cell adhesion and migration programs and metabolic programs compared with 2D cultures^14–17^. Triple-negative breast cancer (TNBC) cells show enhanced mitochondrial metabolism, whereas basal breast cancer cells exhibit increased expression of ECM-interaction genes in 3D microenvironments^16^, highlighting the profound influence of the microenvironment on tumor cell phenotype.

These microenvironment-dependent changes are accompanied by altered therapeutic responses. Numerous studies have shown that cancer cells cultured as multicellular spheroids are generally less sensitive to anticancer agents than cells grown in 2D monolayers^17, 18^, owing in part to diffusion barriers, cell–cell interactions, and extracellular matrix-mediated limitations on drug penetration^19, 20^. Drug responses also vary spatially within spheroids, with cells located in the spheroid core exhibiting distinct sensitivities to chemotherapeutic agents such as paclitaxel (Taxol) and cisplatin compared with cells at the spheroid periphery^21^. Reynolds *et al.* further demonstrated that breast cancer cells embedded diffusely within a 3D ECM exhibit drug responses more closely resembling those of cells at the spheroid periphery, suggesting that diffusely embedded 3D cultures better model invasive or migratory tumor cell populations than conventional spheroids^22^.

Despite these advances, important challenges remain. Most comparisons between 2D and 3D drug responses have focused on spheroid models, where differences in proliferation rates between culture systems can confound interpretation of drug sensitivity^23, 24^. Moreover, although spheroids more closely recapitulate the physiological tumor microenvironment than 2D cell cultures, they are not readily compatible with conventional high-throughput assays for measuring cytotoxicity or gene and protein expression and often require substantial protocol optimization^25, 26^. Consequently, there remains a need for scalable 3D culture platforms that accurately model diffusely growing cancer cells while enabling quantitative, high-throughput assessment of drug responses.

In this study, we developed VITAL, a high-throughput imaging platform for quantitative analysis of drug responses in both 2D and 3D culture systems. By combining volumetric imaging with automated single-cell analysis, VITAL enables direct and precise enumeration of live and dead cells throughout three-dimensional extracellular matrix (ECM) cultures, providing a physiologically relevant platform for studying diffusely embedded cancer cells that model invasive or dormant states^22, 27^. Unlike conventional metabolic assays, such as MTT and ATP-based methods, which infer cell viability from metabolic activity and can be influenced by drug-induced metabolic changes or assay interference^28–30^, VITAL directly quantifies both live and dead cells at single-cell resolution. Furthermore, unlike conventional spheroid assays that rely on indirect measurements such as spheroid diameter, fluorescence intensity, or bulk absorbance, VITAL provides direct volumetric cell counts across the entire 3D culture^31, 32^.

We applied this platform to evaluate the responses of breast cancer cell lines to a panel of anticancer agents and systematically compared drug responses between 2D and 3D microenvironments using both conventional and growth-based response metrics. Together, these studies establish VITAL as a scalable and quantitative platform for evaluating drug efficacy in physiologically relevant tumor models and provide a framework for improving the predictive value of preclinical drug screening.

## Results

### Volumetric live/dead quantification in 3D microenvironments

We established an image-based workflow to directly quantify cell number within 3D extracellular matrix (ECM) environments (**Fig. 1**), termed VITAL (Volumetric Imaging-based Toxicity and Live analysis). To enable efficient live/dead discrimination in a 96-well format, cells were stained with Hoechst 33342 (H33342) and propidium iodide (PI) to label total nuclei and membrane-compromised (dead) cells, respectively. Imaging was performed using a 2× objective, allowing capture of the entire well of a 96 well plate within a single field of view while maintaining sufficient spatial resolution to distinguish individual nuclei. For 3D ECM cultures, volumetric z-stack imaging was performed to encompass the full thickness of the gel, enabling quantification of total cell number throughout the matrix. We developed an image analysis pipeline to accurately detect nuclei and classify live and dead cells based on H33342 and PI fluorescence intensities (**Fig. S1**).

**Figure 1.**
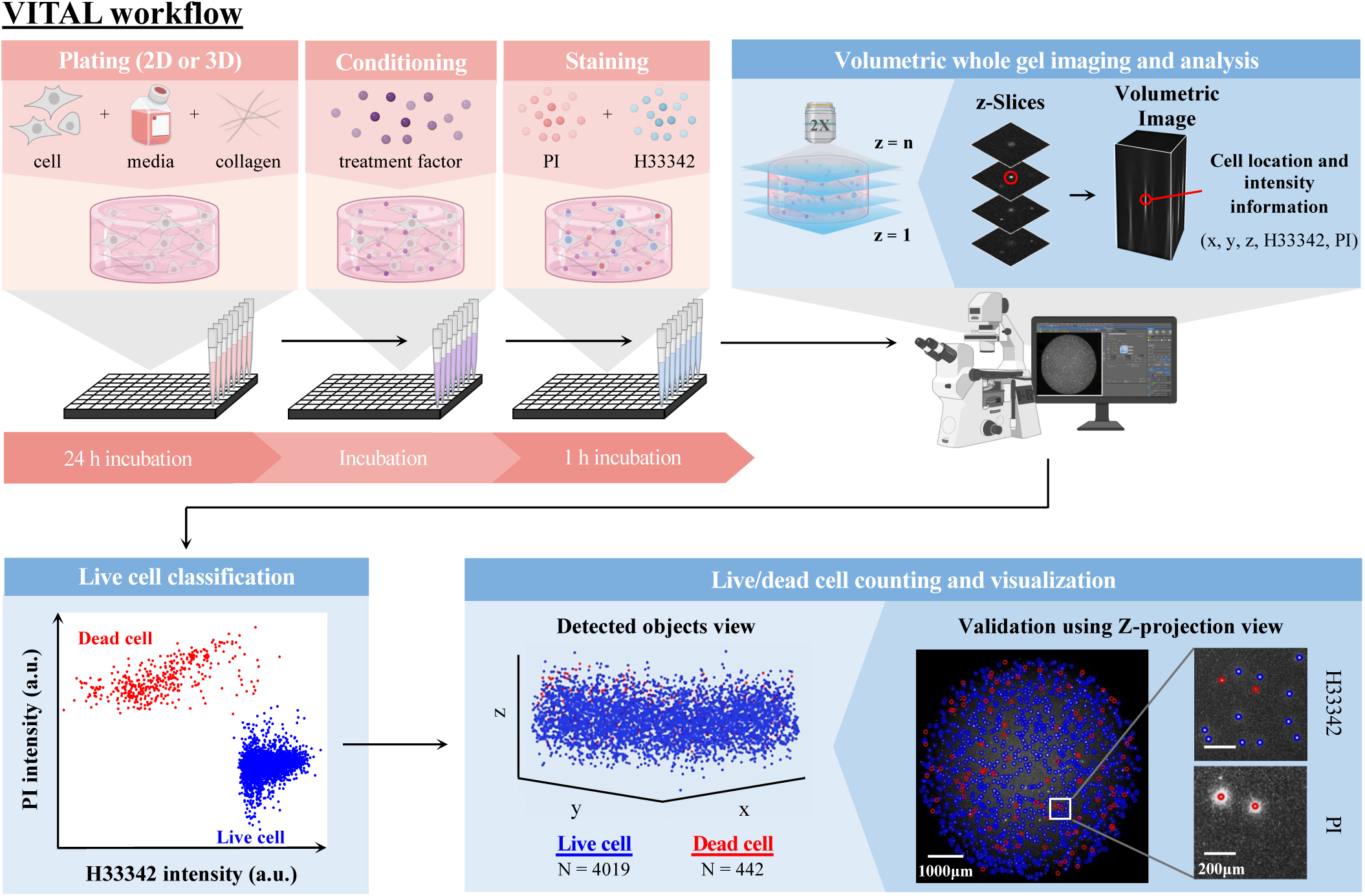
Workflow for cell plating and treatment, staining, volumetric imaging, and automated live/dead cell analysis using VITAL. Breast cancer cell lines were seeded in 96-well plates in both 2D and 3D microenvironments. Following incubation, treatment conditions were applied to each well accordingly. The plates were then stained using Hoechst 33342 (H33342, all cells) and propidium iodide (PI; dead cells). The stained plates were then imaged with an inverted fluorescence microscope with a 2x objective. A single z-position was taken for 2D experiments and approximately 50 z-positions at 50-µm intervals were taken for 3D experiments to encompass the entire gel volume. Cells were identified in the fluorescent images and classified as live or dead based on H33342 and PI intensity.

This workflow enables high-throughput monitoring of cell growth under diverse experimental conditions in 3D ECM systems using a 96-well plate format. The complete workflow—including staining, imaging, and automated quantification—requires less than 2 h to process 96 samples. The same platform can be applied to 2D monolayer cultures. We demonstrate that VITAL achieves >95% counting accuracy for up to 30,000 cells per well in 2D and 15,000 cells per well in 3D cultures within a 96-well format (**Fig. S2**).

### Comparison of the cell growth pattern in 2D and 3D cultures

We first examined the growth dynamics of breast cancer cells embedded in 3D ECM matrices compared to conventional 2D monolayer cultures using VITAL over a 5-day period. A panel of breast cancer cell lines representing diverse receptor subtypes was analyzed, including MCF-7, HCC1954, MDA-MB-231, SUM-159, and SUM-149 (**Fig. 2** and **Table 1**). These cell lines span distinct breast cancer receptor profiles and molecular subtypes. As expected, in 2D culture, most cell lines exhibited an initial exponential growth phase during the first 2–3 days (48–72 h), followed by a deceleration in proliferation as they approached confluence. An exception was HCC1954, which displayed a slower and more linear growth pattern. In contrast to 2D cultures, cells embedded in 3D collagen matrices generally proliferated at substantially lower rates but maintained exponential growth through day 5 (120 h). Notably, HER2⁺ HCC1954 cells failed to persist in 3D collagen matrices, with a progressive decline in live cell number over the 5-day period (**Fig. 2a–f and Fig. S3**). Among all tested cell lines, TNBC SUM-149 cells exhibited the fastest growth rate in 2D culture, with a doubling time of approximately 9 h, whereas HCC1954 showed the slowest growth, with a doubling time of approximately 33 h. ER⁺/HER2⁺ MCF-7 cells exhibited the greatest reduction in proliferation upon transitioning from 2D to 3D culture, with the doubling time increasing from 12.6 h in 2D to 55.6 h in 3D (**Fig. 2g,h**; **Table 1**). While the growth of most cell lines in 2D eventually reached a plateau, SUM-159 cells continued exponential proliferation throughout the 5-day period (**Fig. S3**).

**Figure 2.**
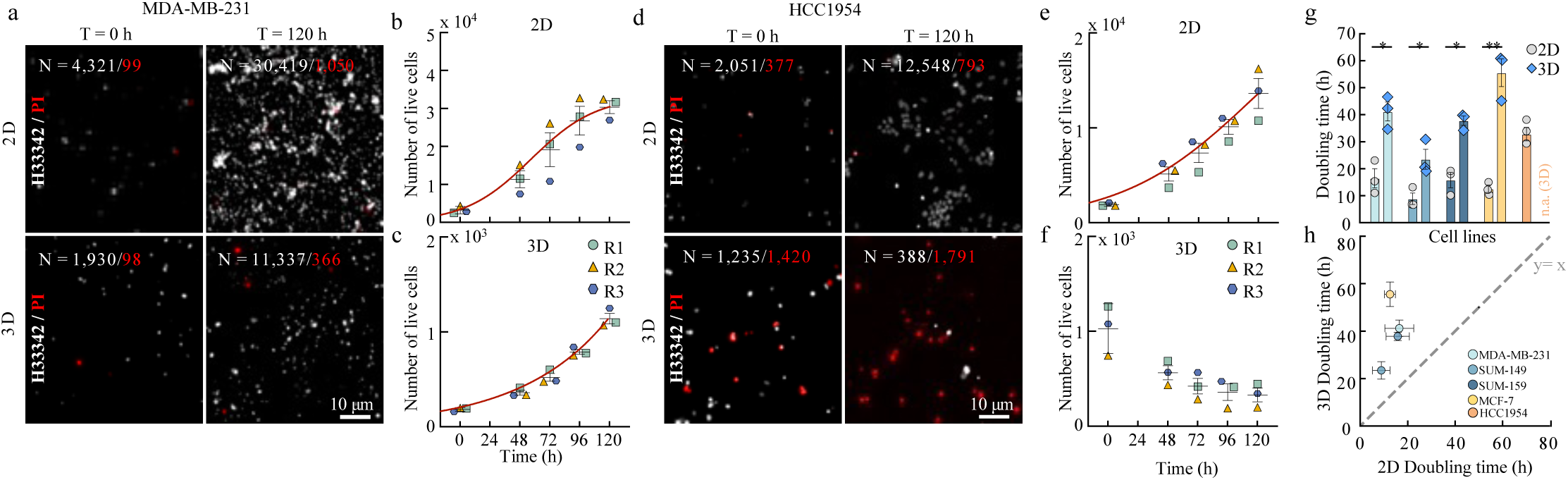
Breast cancer cell lines proliferate more slowly in 3D compared to 2D microenvironments. **(a**) Representative fluorescent images of live MDA-MB-231 cells (nuclei, white) and dead cells (red). (**b-c**) Live cell counts of MDA-MB-231 cultured in 2D (**b**) and 3D (**c**) at 0, 48, 72, 96, and 120 h. (**d**) Representative fluorescent images of live HCC1954 cells (nuclei, white) and dead cells (red). (**e-f**) Live cell counts of HCC1954 cultured in 2D (**e**) and 3D (**f**) at 0, 48, 72, 96, and 120h. (**g**) Doubling times of breast cancer cell lines cultured in 2D and 3D, HCC1954 showed a declining cell population, hence the doubling time is not available in 3D (n.a.). (**h**) Correlation of doubling times between 2D and 3D across cell lines. (**g)** Statistical significance was assessed using two-tailed Student’s *t*-tests. (** p ≤ 0.01, * p ≤ 0.05), Data points represent the mean ± s.e.m. from tee independent biological replicates (n = 3).

**Table 1.** Summary of cell lines used in this study.

| Cell Line | ER | PR | HER2 | Subtype | Disease | BRCA1<br>Mutation | Growth Rate (k) |  | Doubling time (h) |  |
| --- | --- | --- | --- | --- | --- | --- | --- | --- | --- | --- |
|  |  |  |  |  |  |  | 2D | 3D | 2D | 3D |
| MCF-7 | + | + | - | Luminal A | Adenocarcinoma | WT | 0.046 ± 0.009 | 0.017 ± 0.002 | 12.6 ± 1.3 | 55.6 ± 5.1 |
| HCC1954 | - | - | ++ | HER2-positive | Ductal carcinoma | WT | 0.021 ± 0.01 | N/A | 32.8 ± 2.5 | N/A |
| MDA-MB-231 | - | - | - | Triple-negative | Adenocarcinoma | WT | 0.046 ± 0.009 | 0.031 ± 0.004 | 16.5 ± 3.5 | 41.2 ± 3.5 |
| SUM-159 | - | - | - | Triple-negative | Anaplastic Carcinoma | WT | 0.048 ± 0.010 | 0.018 ± 0.001 | 15.8 ± 2.8 | 37.8 ± 1.7 |
| SUM-149 | - | - | - | Triple-negative | inflammatory ductal carcinoma | 2288delT | 0.086 ± 0.017 | 0.031 ± 0.004 | 8.9 ± 2.1 | 23.5 ± 3.7 |

Together, these findings indicate that 3D ECM culture does not uniformly support proliferation across all breast cancer cell types. However, the slower growth kinetics observed in 3D ECM systems provide a valuable framework for evaluating long-term cellular responses to therapeutic treatment.

### Cell responses to taxol in 3D vs. 2D

Taxol is a widely used chemotherapeutic agent for the treatment of breast cancer, particularly in triple-negative and metastatic disease^15^. Taxol stabilizes microtubules and suppresses microtubule dynamics, thereby disrupting normal mitotic progression and ultimately inducing mitotic arrest and cell death^33^. To compare taxol responses between conventional 2D monolayers and 3D ECM cultures, cells were treated with taxol over a concentration range of 0.002 nM to 20 µM or vehicle (DMSO), and cell growth was monitored for five days (120h) using VITAL (**Fig. 3a–c; Fig. S4**). Drug responses were expressed as relative cell number, defined as the percentage of cells remaining relative to the corresponding drug-free control.

**Figure 3.**
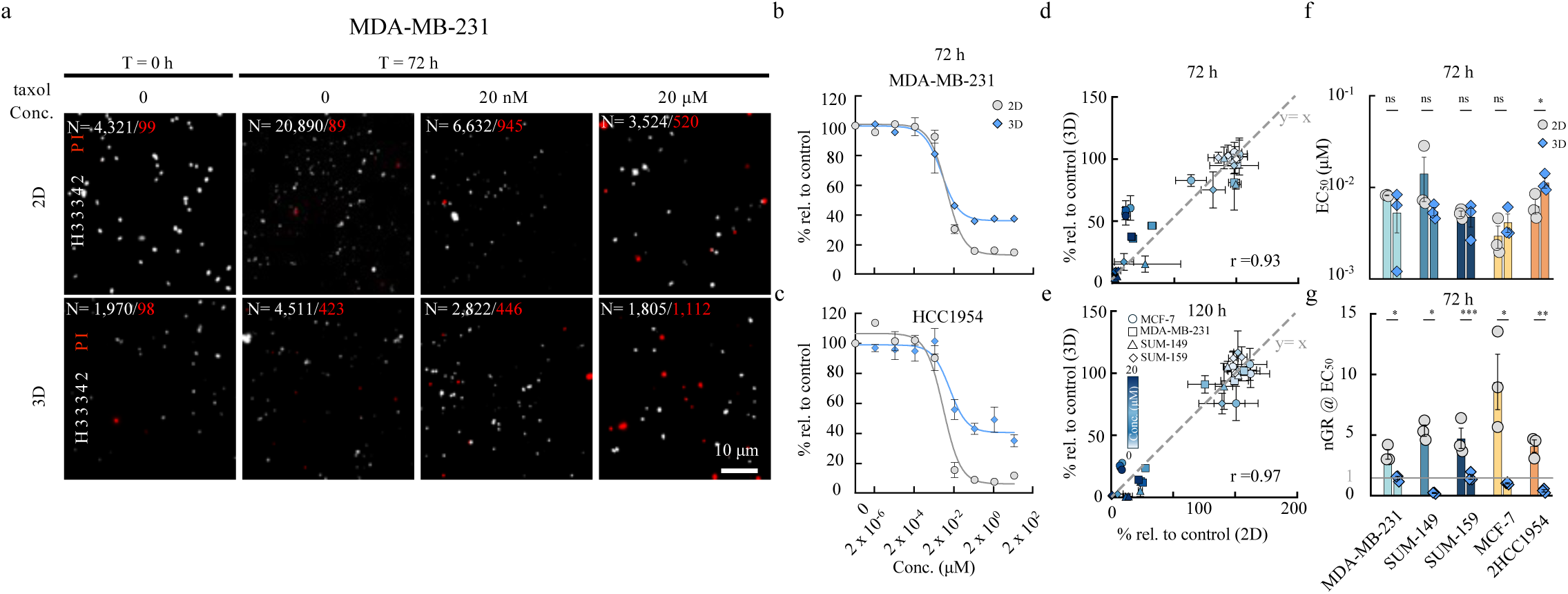
Cellular responses to taxol treatment in 2D and 3D. (**a**) Representative images of live (white) and dead (red) MDA-MB-231 at 0 or 72 h following taxol treatment at 0, 20 nM and 20 μM. (**b-c**) Relative cell count for MDA-MB-231 (**b**) and HCC1954 (**c**) following 72 h taxol treatment in 2D and 3D. (**d-e**) Relative responses to control (%) measured in 2D cultures were compared with those obtained in 3D cultures at 72 h (**d**) and 120 h (**e**) across four cell lines: MCF-7, MDA-MB-231, SUM-149, and SUM-159. The dashed diagonal line represents equal responses in 2D and 3D cultures (y = x). (**f**) Comparison of EC_50_ in 2D and 3D across breast cancer cell lines. (**g**) Normalized growth ratio (nGR) at the respective EC_50_ concentrations after 72 h of treatment in 2D and 3D cultures. **f,g** Statistical significance was assessed using two-tailed Student’s *t*-tests. (*** p ≤ 0.001, ** p ≤ 0.01, * p ≤ 0.05, ns, not significant); Data points represent the mean ± s.e.m. from three independent biological replicates (n = 3).

To assess whether the overall magnitude of cell response to taxol was conserved between culture formats, relative cell numbers were compared between 2D and 3D cultures across the five breast cancer cell lines. A strong positive correlation was observed at both 72 h (Pearson’s *r* = 0.93) and 120 h (*r* = 0.97), indicating that the overall range of taxol responses relative to untreated controls was highly concordant between 2D and 3D cell culture systems (**Fig. 3d,e, Fig. S5**). We next calculated EC₅₀ values from the corresponding dose–response curves. EC₅₀ values tended to be slightly higher in 2D cultures, although the overall difference between 2D and 3D conditions was not statistically significant (**Fig. 3f, Fig. S6**). HCC1954 was the only cell line to exhibit a significant difference, with a higher EC₅₀ in 3D than in 2D (**Fig. 3f**). This difference may in part reflect the reduced baseline growth of HCC1954 cells under drug-free 3D conditions, in which cell number declined over time even in the absence of taxol. Together, these results demonstrate that both the overall magnitude of taxol response and EC₅₀ measurements were broadly concordant between 2D and 3D cultures, while revealing cell line-specific differences that may be influenced by baseline growth behavior.

However, EC₅₀ alone does not fully capture the magnitude or biological nature of the drug response^34, 35^. For example, MDA-MB-231 cells exhibited comparable EC₅₀ values in 2D and 3D cultures (8.20 and 5.31 nM, respectively), despite a marked difference in maximal response. At the highest taxol concentration, the relative cell population was approximately 10% in 2D cultures compared with approximately 40% in 3D cultures (**Fig. 3b, Fig. S4**). Thus, similar EC₅₀ values can be associated with substantially different treatment outcomes.

To further characterize these differences, we introduced the normalized growth ratio (nGR), defined as the ratio of cell number at the end of treatment to that at treatment initiation, nGR = Nₜ/N₀. An nGR > 1 indicates net population expansion, an nGR ≈ 1 indicates growth arrest, and an nGR < 1 indicates net cell loss relative to the initial population. Across the tested cell lines, treatment at the respective EC₅₀ concentrations generally resulted in continued population expansion in 2D cultures (nGR > 1), whereas the corresponding 3D cultures exhibited markedly reduced growth or net cell loss (nGR ≤ 1) (**Fig. 3g**). For example, SUM149 cells increased 5.4-fold relative to the initial cell number in 2D cultures but declined to 20% of the initial population in 3D at their respective EC₅₀ concentrations. Similarly, MCF-7 cells expanded 9.3-fold in 2D but remained essentially growth arrested in 3D (nGR = 1.01).

Together, these findings demonstrate that comparable EC₅₀ values between 2D and 3D cultures do not necessarily reflect comparable biological responses. Incorporating growth-based measurements such as nGR provides complementary information by distinguishing continued proliferation, growth arrest, and net population loss that are not captured by EC₅₀ alone.

### Growth-based metrics reveal drug responses beyond EC₅₀

Since EC₅₀ provides only a single endpoint measure of drug potency, we next evaluated temporal growth responses using the normalized growth ratio (nGR) across a range of paclitaxel concentrations in both 2D and 3D cultures (**Fig. 4a**). The nGR metric characterizes cellular responses as continued proliferation (nGR > 1), growth arrest (nGR ≈ 1), or net cell loss (nGR < 1), thereby capturing dynamic changes in population growth over time. Across all cell lines, paclitaxel began suppressing proliferation at similar concentration (∼0.02 µM) in both 2D and 3D cultures, as indicated by a marked reduction in nGR (**Fig. 4a**). However, cellular responses diverged substantially at higher concentrations and longer treatment durations. MCF-7 cells largely underwent growth arrest (nGR ≈ 1), whereas the TNBC cell lines and HCC1954 exhibited net cell loss (nGR < 1) following prolonged treatment. These differences were not reflected by EC₅₀ values, which remained relatively similar across cell lines (**Fig. 3d–f; Fig. S6**)^36, 37^.

**Figure 4.**
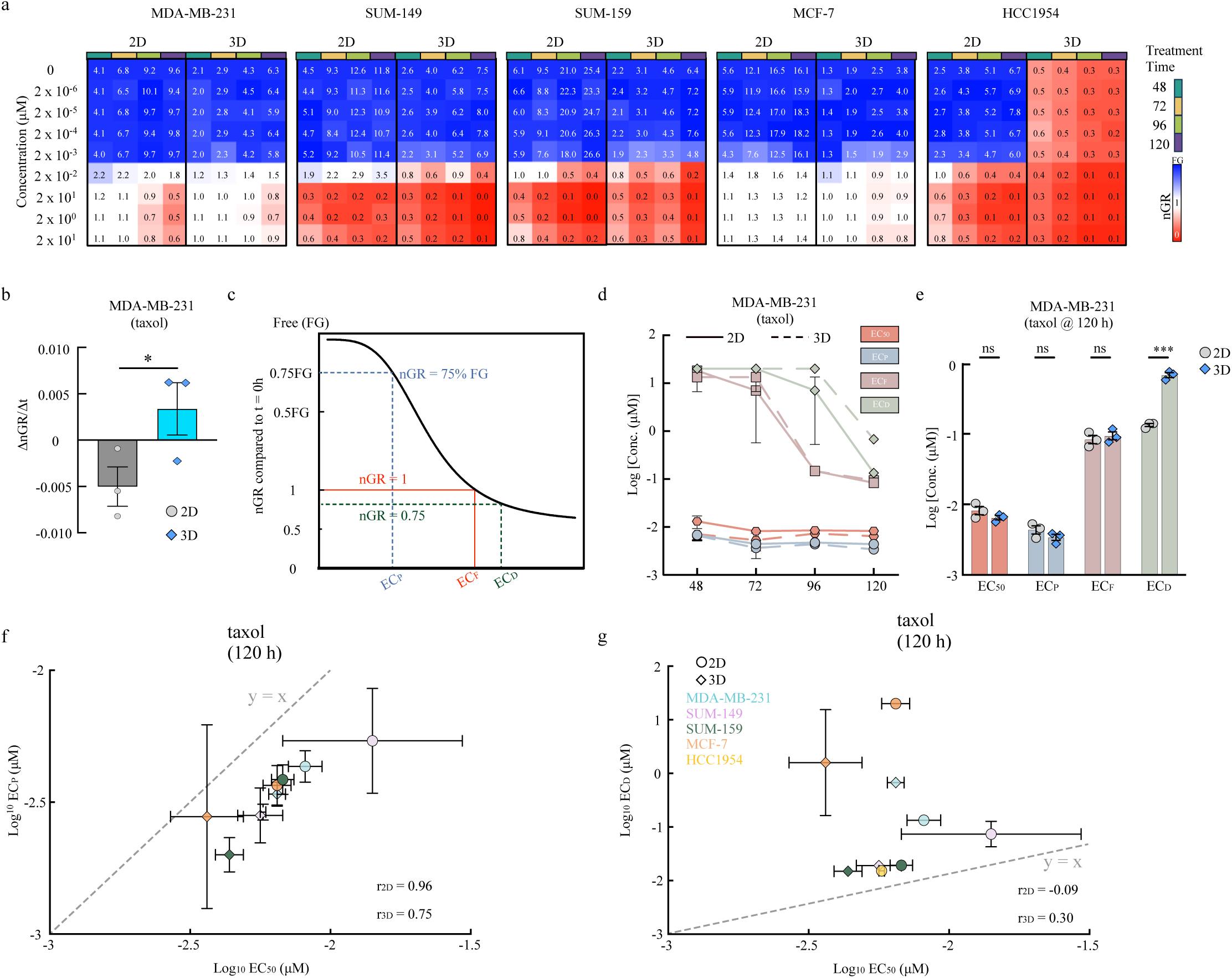
Comparison of drug responses between 2D and 3D cultures. (**a**) Heatmaps showing the nGR across drug concentrations and time points (48, 72, 96, and 120 h) for five breast cancer cell lines (MCF-7, MDA-MB-231, SUM-149, SUM-159, HCC1954) in 2D and 3D. Colors indicate growth states: blue, uninhibited growth; white, partial growth inhibition; red, net cell loss. (**b**) Comparison of the temporal slopes of nGR for MDA-MB-231 treated with 2 × 10⁻² μM taxol in 2D and 3D cultures, data are shown as mean ± s.e.m. (**c**) Schematic illustration of nGR-based dose-response curves. The blue dashed line indicates effective concentration for partial inhibition (EC_P_; 75% free growth, which is untreated condition), The red dashed line indicates effective concentration for full growth arrest (EC_F_, stalled growth, nGR ≈ 1), and the green dashed line indicates the effective concentration for death induction (EC_D_; net cell loss, nGR < 1). (**d**) Temporal drug dynamics of EC_50_, EC_P_, EC_F_, EC_D_ values in MDA-MB-231 in 2D (solid lines) and 3D (dashed lines) cultures across time points under taxol treatment. (**e**) Quantitative comparison of EC_50_, EC_P_, EC_F_, EC_D_ between 2D and 3D cultures of MDA-MB-231 under taxol treatment at 120 h. (**f,g**) Correlations between EC_50_ and EC_P_ (**f**), and between EC_50_ and EC_D_ (**g**) at 120 h for all cell lines treated with taxol in 2D and 3D. The dashed diagonal line represents equal concentration in 2D and 3D cultures (y = x). Each dot denotes a single cell line is shown as mean ± s.e.m with three biological replicates (n = 3). **b,e** Statistical significance was assessed using two-tailed Student’s *t*-tests. (*** p ≤ 0.001, * p ≤ 0.05, ns, not significant).

Comparison of nGR profiles between 2D and 3D cultures further highlighted the limitations of EC₅₀. In MCF-7 cells treated with an effective paclitaxel concentration (∼0.02 µM), nGR exceeded 1 in 2D cultures but remained approximately 1 in 3D cultures, indicating greater growth inhibition in the 3D microenvironment despite a slightly higher EC₅₀ (**Fig. 3g**; **Fig. 4a**). Conversely, MDA-MB-231 cells exhibited a declining growth trend in 2D culture, whereas they continued to proliferate steadily in 3D throughout the 5-day treatment period (**Fig. 4a,b**), demonstrating enhanced survival in the 3D microenvironment that was not apparent from EC₅₀ values alone. Together, these findings demonstrate that drug responses are strongly influenced by the cellular microenvironment and that EC₅₀ alone does not distinguish continued proliferation, growth arrest, and net cell loss.

To quantitatively characterize these distinct response states, we defined three complementary growth-based metrics: the effective concentration for partial inhibition EC_P_, the effective concentration for full growth arrest EC_F_, and the ECD represents the minimum concentration required to induce net cell loss (nGR < 1) (**Fig. 4c**). EC_P_ represents the minimum concentration required to reduce net population growth by 25% relative to untreated controls, EC_F_ denotes the concentration required to achieve complete growth arrest (nGR ≈ 1), and EC_D_ represents the minimum concentration required to induce net cell loss (nGR < 1).

Using MDA-MB-231 cells as an example, EC₅₀, EC_P_, and EC_F_ were comparable between 2D and 3D cultures at all time points, indicating similar thresholds for suppressing proliferation (**Fig. 4d,e**). In contrast, EC_D_ was significantly higher in 3D cultures than in 2D after 120 h (0.68 ± 0.05 µM versus 0.13 ± 0.01 µM; **Fig. 4d,e**; **Supplementary Data 1**), indicating that substantially higher paclitaxel concentrations were required to induce net cell loss within the 3D microenvironment.

Across all cell lines, EC₅₀ closely tracked EC_P_ and was strongly correlated with this metric (Pearson’s *r* = 0.96 in 2D and *r* = 0.75 in 3D at 120 h; **Fig. 4f**), indicating that EC₅₀ primarily reflects the onset of growth inhibition. In contrast, EC_D_ values were consistently higher than EC₅₀ and showed little correlation with EC₅₀ (Pearson’s *r* = −0.09 in 2D and *r* = 0.30 in 3D; **Fig. 4g**), demonstrating that drug potency alone does not predict the concentration required to induce net population loss.

Collectively, these results show that although EC₅₀ values often suggest similar drug responses between 2D and 3D cultures, growth-based metrics reveal substantial differences in proliferative behavior and death induction. Integrating dynamic measurements such as nGR, EC_P_, EC_F_, and EC_D_ therefore provides a more comprehensive and biologically meaningful assessment of therapeutic response in physiologically relevant 3D microenvironments.

The ability of VITAL to simultaneously quantify live and dead cells also enables direct measurement of cell viability, defined as the fraction of live cells within the total cell population, in addition to conventional measurements of cell number normalized to untreated controls (**Fig. 4a; Fig. S4; Fig. S7**). We next examined the relationship between cell viability and normalized growth ratio nGR. Across all treatment time points, nGR showed a strong positive correlation with the live-cell fraction (Pearson’s *r* > 0.8). Notably, conditions in which the live-cell fraction fell below 50% consistently exhibited net population loss (nGR < 1) across all tested cell lines. In contrast, when the live-cell fraction exceeded 50%, nGR values were substantially more variable, ranging from approximately 1, indicating growth arrest, to substantially greater than 1, indicating continued population expansion. Thus, a high viable-cell fraction can correspond to markedly different proliferative states, ranging from growth arrest to substantial population expansion, which cannot be distinguished using viability measurements alone.

Together, these results demonstrate that the simultaneous measurement of cell viability and population growth by VITAL provides complementary information for distinguishing cell death, growth arrest, and continued proliferation following drug treatment (**Fig. S8**).

### Differential drug responses in 2D and 3D breast cancer models

We next systematically compared drug sensitivities between conventional 2D monolayer cultures and 3D ECM microenvironments. Using VITAL, we quantified growth responses to four additional FDA-approved breast cancer therapeutics (**Table 2**) across four breast cancer cell lines cultured under both conditions (**Fig. S9 – Fig. S10**). Overall, EC_P_ and EC_F_ values were strongly correlated between 2D and 3D cultures (Pearson’s *r* > 0.8; **Fig. 5a,b**), indicating that variability in drug potency among different therapeutics exceeded the overall effects of culture format.

**Figure 5.**
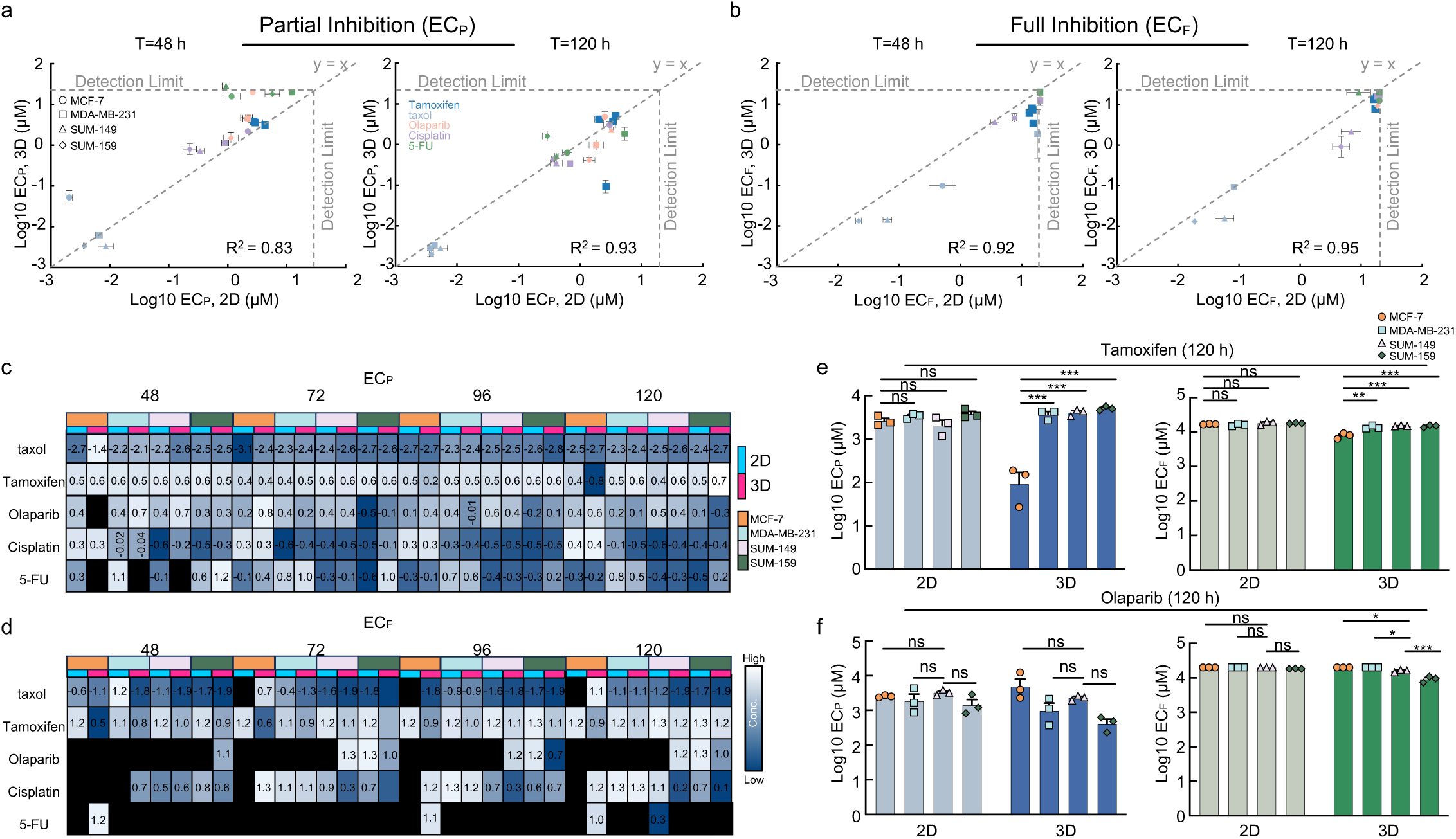
Comparison of cellular responses to FDA-approved breast cancer therapeutics in 2D and 3D cultures. (**a,b**) Correlation analyses of partial (EC_P_; **a**) and full (EC_F_; **b**) inhibitory concentrations for FDA-approved breast cancer drugs measured in 2D and 3D microenvironments at 48 h and 120 h. Vertical and horizontal dashed lines represent the maximum tested drug concentrations in 2D and 3D cultures, respectively. The dashed diagonal line represents equal concentration in 2D and 3D cultures (y = x). Each dot denotes a single compound. Data are shown as mean ± s.e.m. (**c,d**) Heatmaps showing partial (EC_P_; top; **c**) and full (EC_F_; bottom; **d**) inhibitory concentrations for FDA-approved breast cancer drugs across four cell lines (MCF-7, MDA-MB-231, SUM-149, and SUM-159); four time points (48, 72, 96, and 120 h) in 2D and 3D microenvironments. Black box indicates cases in which inhibitory concentration could not be determined within the tested dose range. Values represent the median of three independent biological replicates (n = 3). (**e-f**) Comparison of cellular responses to tamoxifen (**e**), and olaparib (**f**) at 120 h. Each dot represents an independent biological replicate. Statistical significance was assessed using one-way ANOVA with post hoc Tukey’s test. (*** p ≤ 0.001, ** p ≤ 0.01, * p ≤ 0.05, ns, not significant).

**Table 2.** Summary of drugs used in this study.

| Drug | Type | Mechanism of action | Primarily for |
| --- | --- | --- | --- |
| 5-FU | Antimetabolite | Inhibits thymidylate synthase and disrupts DNA synthesis | TNBC |
| Cisplatin | Pt-based antineoplastic | Induces DNA crosslinks and DNA damage | TNBC with BRCA mutation |
| Olaparib | PARP inhibitor | Inhibits PARP-mediated DNA repair | TNBC with BRCA mutation |
| Tamoxifen | Anti-estrogen | Selective estrogen receptor modulator | Hormone receptor positive |
| taxol | Antimitotic | Stabilizes microtubules and disrupts mitotic progression | TNBC |

Despite this global concordance, growth-based metrics revealed clear microenvironment-dependent differences in drug response. Although EC₅₀ values were generally comparable between 2D and 3D cultures, EC_P_ and EC_F_ exposed distinct differences in proliferative behavior (**Fig. S9–S15)**. At 48 h, EC_P_ values were generally higher in 3D cultures, indicating greater resistance to partial growth inhibition. By 120 h, this trend largely reversed, with 2D cultures exhibiting higher EC_P_ values despite similar EC₅₀ values between culture conditions (**Fig. S15**). In contrast, EC_F_ values were consistently higher in 2D than in 3D across most drugs and time points, indicating that complete growth arrest required higher drug concentrations in 2D cultures (**Fig. 5a–d, Fig. S16)**. Notably, EC_F_ values frequently approached the highest tested concentrations after prolonged treatment, indicating that sustained exposure alone was often insufficient to fully suppress proliferation.

To determine which culture model and response metrics best capture biologically expected drug responses, we examined subtype-specific therapies. Tamoxifen, a selective estrogen receptor (ER) modulator, is expected to preferentially inhibit ER-positive breast cancer^38, 39^. Under conventional 2D culture conditions, however, MCF-7 cells showed little distinction from the TNBC cell lines throughout the treatment period. In contrast, under 3D culture conditions, MCF-7 cells exhibited significantly lower EC_P_ and EC_F_ values after 120 h than the non-ER-positive cell lines (**Fig. 5c–e**), consistent with their expected sensitivity to tamoxifen. These findings suggest that combining a physiologically relevant 3D microenvironment with extended treatment duration more accurately recapitulates clinically relevant endocrine responses. A similar pattern was observed for olaparib, a PARP inhibitor approved for BRCA1/2-mutated breast cancers. SUM149 cells, which harbor a pathogenic BRCA1 mutation (c.2288delT), exhibited markedly greater sensitivity under 3D culture conditions after 120 h, with significantly lower EC_F_ values than MDA-MB-231 and MCF-7 cells (**Fig. 5f**). Interestingly, SUM159 cells also demonstrated increased sensitivity to olaparib in 3D despite lacking the same BRCA1 mutation^40^, suggesting that the 3D microenvironment may uncover additional determinants of PARP inhibitor responsiveness beyond canonical BRCA1 deficiency.

Finally, we compared drug response profiles across cell lines for all tested therapeutics. In 2D cultures, MDA-MB-231 responses correlated only weakly with those of the other cell lines, including ER-positive MCF-7, particularly for EC_F_ values. In contrast, under 3D culture conditions, both EC₅₀ and EC_F_ profiles clustered the three TNBC cell lines together while clearly separating them from MCF-7 (**Fig. S18**). These findings indicate that the 3D microenvironment more faithfully preserves subtype-specific drug response relationships among breast cancer models. Collectively, these results demonstrate that although overall drug potency is broadly conserved between 2D and 3D cultures, growth-based response metrics reveal substantial microenvironment-dependent differences in therapeutic response. By more accurately resolving subtype-specific sensitivities and distinguishing partial growth inhibition from complete growth arrest, physiologically relevant 3D cultures combined with growth-based analysis provide a more informative framework for preclinical drug evaluation.

## Discussion

In this study, we developed VITAL, a high-throughput volumetric imaging platform that enables direct single-cell quantification of proliferation and cell death in three-dimensional (3D) extracellular matrix (ECM) cultures. Using this platform, we demonstrated that although conventional EC₅₀ measurements are often similar between 2D and 3D cultures, they can mask substantial differences in growth dynamics and cytotoxic responses. Growth-based metrics, including nGR, EC_P_, EC_F_, and EC_D_, revealed that drug responses are strongly influenced by the cellular microenvironment and distinguished biological outcomes that were not captured by EC₅₀ alone. Furthermore, subtype-specific therapeutic responses, including tamoxifen sensitivity in ER-positive cells and olaparib sensitivity in BRCA1-mutant cells, were more faithfully resolved under 3D culture conditions, highlighting the importance of physiologically relevant models for preclinical drug evaluation.

Our findings are consistent with accumulating evidence that the tumor microenvironment profoundly influences cancer cell behavior and therapeutic response^41–44^. Previous studies have shown that 3D breast cancer models more faithfully preserve tissue architecture, extracellular matrix interactions, diffusion gradients, and microenvironment-dependent signaling than conventional monolayer cultures^41, 43, 45, 46^. These features alter proliferation, metabolism, survival signaling, and drug penetration, thereby reshaping therapeutic sensitivity^44, 47, 48^. Our data extend these observations by demonstrating that many of the apparent differences between 2D and 3D drug responses arise not from changes in EC₅₀ itself, but from fundamentally different growth trajectories following treatment. In particular, we found that concentrations required to induce complete growth arrest or net cell loss frequently diverged between 2D and 3D cultures despite similar EC₅₀ values, indicating that drug potency alone does not adequately describe therapeutic outcome in structurally complex microenvironments.

These findings also highlight an important limitation of conventional endpoint-based drug screening. EC₅₀ primarily reflects the concentration required to reduce population growth relative to untreated controls, but does not distinguish continued proliferation, growth arrest, or net cell loss. Consequently, two experimental conditions with similar EC₅₀ values may exhibit markedly different biological responses, particularly when baseline proliferation rates differ. By integrating growth-based response metrics with volumetric single-cell imaging, VITAL enables quantitative discrimination between partial growth inhibition, cytostasis, and cytotoxicity, providing a more comprehensive assessment of drug efficacy across distinct microenvironments.

Compared with conventional metabolic assays such as MTT, MTS, or ATP-based measurements, VITAL directly enumerates live and dead cells rather than inferring viability from metabolic activity. Because metabolic assays can be influenced by changes in mitochondrial activity, cellular metabolism, or assay interference, they may overestimate or underestimate viable cell number and antiproliferative effects^28–30^. Likewise, proliferation assays such as BrdU or EdU labeling measure DNA synthesis during a limited labeling period but do not quantify cumulative population growth or cell loss over time^49, 50^. By directly measuring changes in cell number relative to the initial seeding density, VITAL provides a dynamic and quantitative measure of growth that is readily applicable to both 2D monolayer and diffusely embedded 3D cultures. Unlike spheroid-based approaches that rely on indirect measurements such as spheroid diameter or bulk fluorescence intensity, volumetric single-cell imaging enables direct quantification throughout the entire ECM volume, making the platform well suited for high-throughput analysis of diffusely growing tumor cells.

Several limitations should be considered. First, our study was performed using established breast cancer cell lines embedded in collagen matrices, which do not fully recapitulate the cellular and extracellular complexity of human tumors. Second, although growth-based metrics distinguished cytostatic and cytotoxic responses, the molecular mechanisms underlying the observed microenvironment-dependent differences remain to be determined. Future studies integrating transcriptomic or proteomic profiling with patient-derived organoids, immune or stromal components, and *in vivo* validation will help define how extracellular matrix interactions regulate therapeutic sensitivity and resistance.

In conclusion, our study demonstrates that growth-based single-cell analysis provides substantially greater biological insight than conventional endpoint measurements for evaluating anticancer drug responses. By combining volumetric imaging with quantitative growth metrics in a scalable 96-well format, VITAL offers a practical platform for physiologically relevant preclinical drug screening and provides a framework for improving the biological relevance of *in vitro* therapeutic evaluation.

## Acknowledgements

The authors acknowledge the following sources of support: UG3CA275681(PHW), UH3CA275681(PHW); U54AR081774 (DW); U54CA268083 (DW); R01CA300052 (DW). We gratefully acknowledge the Daniele Gilkes laboratory for providing the SUM149 and SUM159 breast cancer cell lines used in this study.

## Author contributions

C.J., Q.Z., and P.H.W. designed the experiments. C.J. and T.C. collected the data. Q.Z. and P.H.W. developed the analytical tools. C.J. and T.C. analyzed the data. C.J., T.C., and P.H.W. prepared the figures and supplementary materials. C.J., P.H.W., and D.W. wrote the manuscript. D.W. and P.H.W. edited the manuscript.

## Competing interests

All authors declare no competing interests.

## Materials and Methods

### Cell culture

Human breast cancer cell lines MDA-MB-231, MCF-7 and HCC-1954 were obtained from ATCC. SUM149 and SUM159 cells were obtained from a collaborator, as previously described^51^. All cell lines were cultured in high-glucose DMEM (4.5 g/L glucose, Corning) containing 10% fetal bovine serum (FBS, Corning) and 1% penicillin-streptomycin (Gibco). Cells were maintained at 37 °C and 5% CO_2_ and sustained in culture between passages 3 and 10.

### 2D and 3D growth and cytotoxicity assays

Breast cancer cells were cultured until they reached approximately 80% confluency, at which point they were trypsinized and collected for cell counting using a hemocytometer. For 2D growth and cytotoxicity assays, cells were seeded into 96-well black-wall flat-bottom plates at densities of 2,000 cells/well for MDA-MB-231, SUM-149, and HCC1954, and 1,000 cells/well for SUM-159 and MCF-7, due to their higher proliferation rates. For 3D growth and cytotoxicity assays, suspended cells were mixed with high-glucose DMEM and high-concentration collagen type I (Corning) to a final collagen concentration of 2.5 mg/mL. This mixture was prepared on ice in a transfer well (Thermosphere) and incubated on ice for 30 minutes. Then, 100 μL of the cell–collagen mixture was dispensed into each well of a 96-well black-wall flat-bottom plate placed on a 37°C hotplate. Plates were transferred to a cell culture incubator for 30 minutes to allow gel solidification, after which 100 μL of cell culture medium was added to each well. Cells were incubated overnight before compound treatment. Testing compounds were serially diluted from stock solutions to the desired concentrations. For treatment, 25 μL of compound solution mixed with 25 μL of culture medium was added to 2D cultures, while 50 μL of compound solution mixed with 50 μL of culture medium was added to 3D cultures, resulting in the final treatment concentrations. The compounds tested included 5-fluorouracil (5-FU; Catalog No. S1209), cisplatin (Catalog No. S1166), paclitaxel (Taxol; Catalog No. S1150), and tamoxifen (Catalog No. S1238), which were obtained from Selleckchem. Olaparib (Catalog No. SML3705) was obtained from Merck. All drugs were prepared as 10 mM stock solutions in DMSO or deionized water, as appropriate, and stored at −80 °C.

### Live/dead staining and imaging

Cells cultured in 2D monolayers or 3D extracellular matrix (ECM) gels in 96-well plates were stained simultaneously with Hoechst 33342 (H33342; Thermo Fisher Scientific, Cat. No. 62249) at a final concentration of 1 μM and propidium iodide (PI; Thermo Fisher Scientific, Cat. No. P1304MP) at a final concentration of 500 nM. The staining solution was added directly to each well without aspirating the culture medium to minimize disturbance of the cell monolayer or ECM gel. A total of 25 μL and 50 μL of the H33342/PI staining solution was added to each well for 2D and 3D cultures, respectively, followed by incubation for 1 h. Following staining, 96-well plates were transferred to a Nikon Eclipse Ti-E inverted fluorescence microscope (Nikon Instruments) equipped with an on-stage incubator (Tokai Hit, INU-TIZW), which maintained physiological culture conditions at 37°C and 5% CO₂ throughout image acquisition. Images were acquired using a 2× objective. For both 2D and 3D cultures, the field of view was adjusted to capture the entire well within a single image. In 2D cultures, images were acquired at the focal plane of the cell monolayer. For 3D ECM cultures, the upper and lower boundaries of the gel were identified manually, and volumetric z-stack images were acquired from the bottom to the top of the gel at 50-μm intervals, yielding approximately 50 optical sections per well. This imaging strategy enabled complete volumetric visualization of the embedded cell population for subsequent automated single-cell quantification.

### Image processing and cell counting

Fluorescence images were analyzed using a custom MATLAB pipeline (MathWorks, Natick, MA, USA) developed for automated single-cell detection and quantification in 2D monolayer and 3D extracellular matrix cultures. Hoechst 33342 and PI images were acquired as channel 1 and channel 2, respectively. For visualization of 3D datasets, maximum-intensity projections of both fluorescence channels were generated from the complete z-stack. To correct for uneven illumination and background fluorescence, each image was downsampled, median filtered, and resized to its original dimensions to generate a background estimate, which was then subtracted from the raw image. Background-corrected images were subsequently band-pass filtered to suppress high-frequency noise and enhance nuclear features^52^. Candidate nuclei were detected independently in the H33342 and PI channels using local intensity maxima and predefined intensity thresholds. The centroid coordinates and fluorescence intensities of detected objects were refined using intensity-weighted centroid localization. For 2D cultures, analysis was performed on a single focal plane corresponding to the cell monolayer. To prevent duplicate counting of nuclei labeled by both H33342 and PI, PI-positive objects located within 5 pixels of a H33342-positive object were considered to represent the same cell and were merged into a single detection. For each detected cell, the x- and y-coordinates and fluorescence intensities in both channels were recorded. H33342-positive, PI-negative nuclei were classified as live cells, whereas PI-positive nuclei were classified as dead cells. For 3D cultures, nuclei were first detected independently in each optical section. PI-positive objects located within 3 pixels of a H33342-positive object in the same z-plane were merged to avoid duplicate detection. Corresponding nuclear detections were then linked across adjacent z-sections using a nearest-neighbor tracking algorithm to reconstruct individual 3D objects^11, 12^. For each reconstructed nucleus, the x- and y-coordinates were assigned from the optical section containing the maximum nuclear fluorescence intensity, whereas the z-coordinate was calculated as an intensity-weighted centroid across the linked sections. The final output for each cell included its 3D coordinates, H33342 and PI fluorescence intensities, object identifier, and the number of optical sections occupied by the reconstructed nucleus. Live and dead cells were classified according to their H33342 and PI fluorescence profiles, and total live and dead cell numbers were calculated for each well. Automated cell counts and positional information were exported for subsequent analyses of cell proliferation, viability, normalized growth ratio, and drug-response metrics.

### Statistical analysis

Statistical analyses were performed using GraphPad Prism 10 (GraphPad Software, Inc.). Data are presented as mean ± standard error of the mean (s.e.m.), unless otherwise stated. Biological replicate numbers are indicated in the corresponding figure legends. Comparisons between two groups were performed using unpaired two-tailed Student’s t-tests. Comparisons among multiple groups were performed using one-way analysis of variance (ANOVA), followed by appropriate post hoc multiple-comparison tests where applicable. Statistical significance was defined as P < 0.05. Significance levels are indicated as follows: *P < 0.05, **P < 0.01, and ***P < 0.001.

### Code availability

The source code is available on GitHub at https://github.com/pixel-bio

## Notes

### Competing Interest Statement

The authors have declared no competing interest.

