## supplementary figures for "VITAL-3D: Volumetric Single-Cell Quantification Reveals Microenvironment-Dependent Drug Responses in Breast Cancer"

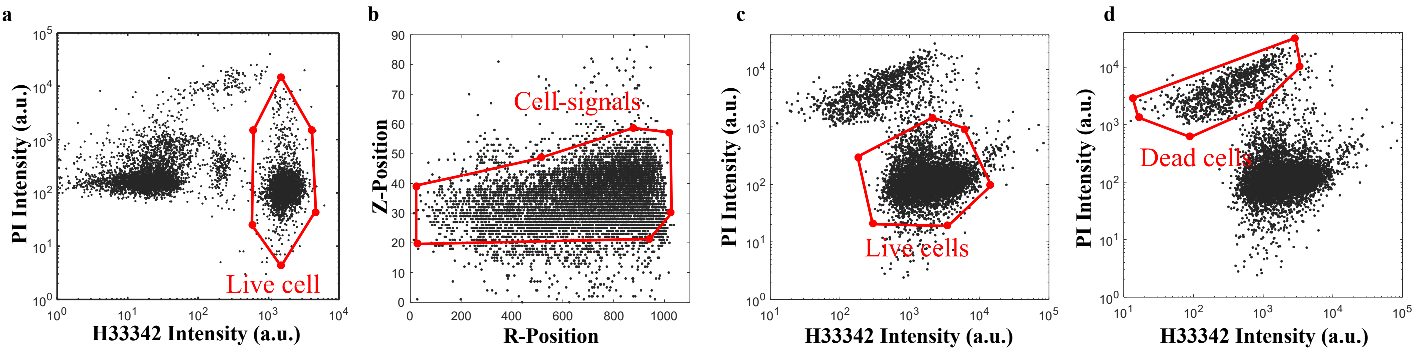


**Figure S1. Noise-reduction and cell-selection gating strategies for 2D (a) and 3D (b-d).** Live and dead cells were identified using manual gating strategies which were applied following automated detection. Potential noise, debris, and noncellular signals were excluded using these gating criteria. **(a)** Gating for live cells using H33342 and PI intensity. **(b)** Removal of noise and debris via positive gating for possible cell locations using Z-position, H33342 intensity, and radial (R) position of the detected signal. **(c)** Gating for live cells using H33342 and PI intensity. **(d)** Gating for dead cells using H33342 and PI intensity.

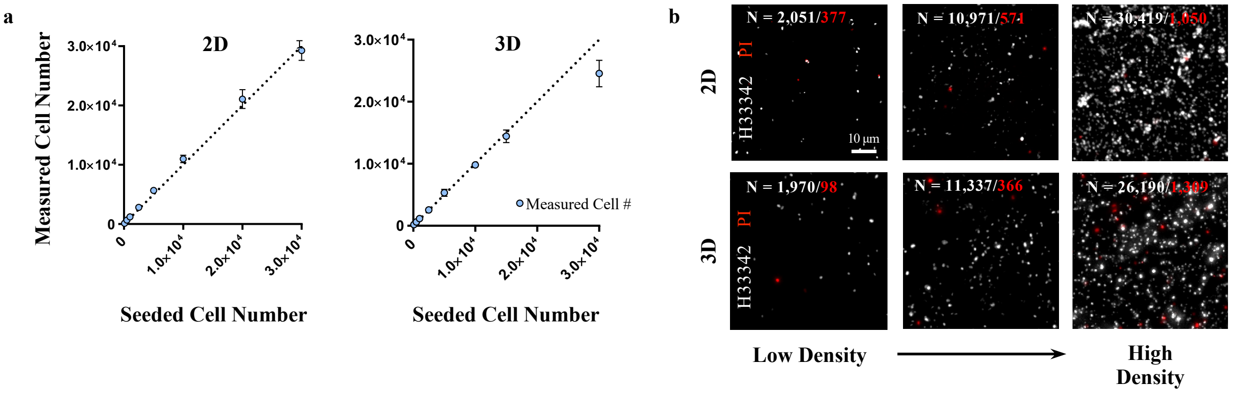


**Figure S2. Validation of workflow and automated live-dead cell analysis.** MDA-MB-231 were seeded at densities ranging from 100 to 30,000 cells/well in 96-well plates under 2D and 3D culture conditions. To verify the accuracy of our live-dead cell analysis platform, the measured cell count 24 hours following plating was compared to the seeded cell number. Data points represent the mean ± s.e.m. from three independent biological replicates (n = 3).

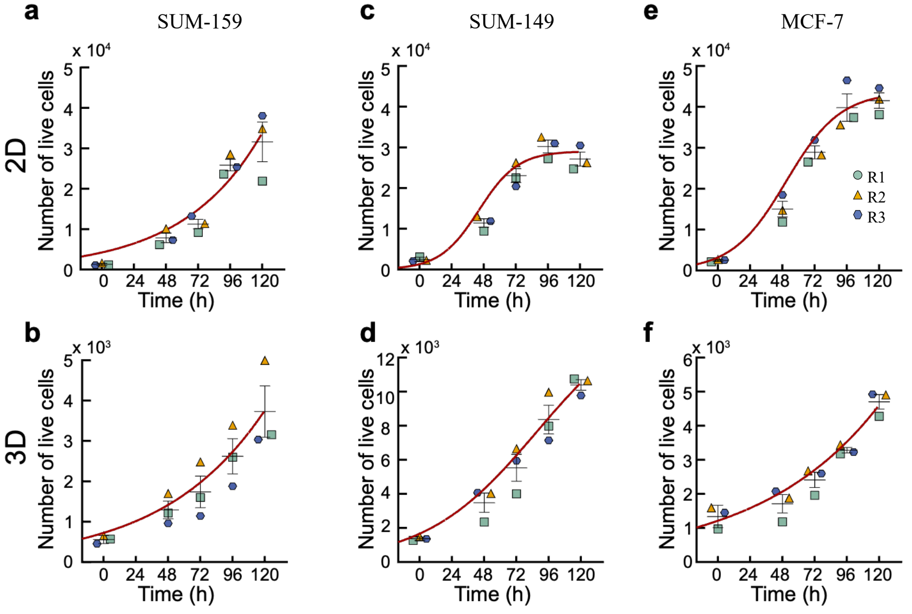


**Figure S3. Growth kinetics of SUM-159, SUM-149, and MCF-7 cell lines in 2D and 3D microenvironments.** Live cell count of SUM-159, SUM-149 and MCF-7 at 0, 48, 72, 96, 120 h culturing in 2D (**a, c, e**), and 3D (**b, d, f**) microenvironments. Data points represent the mean ± s.e.m. from three independent biological replicates (n = 3).

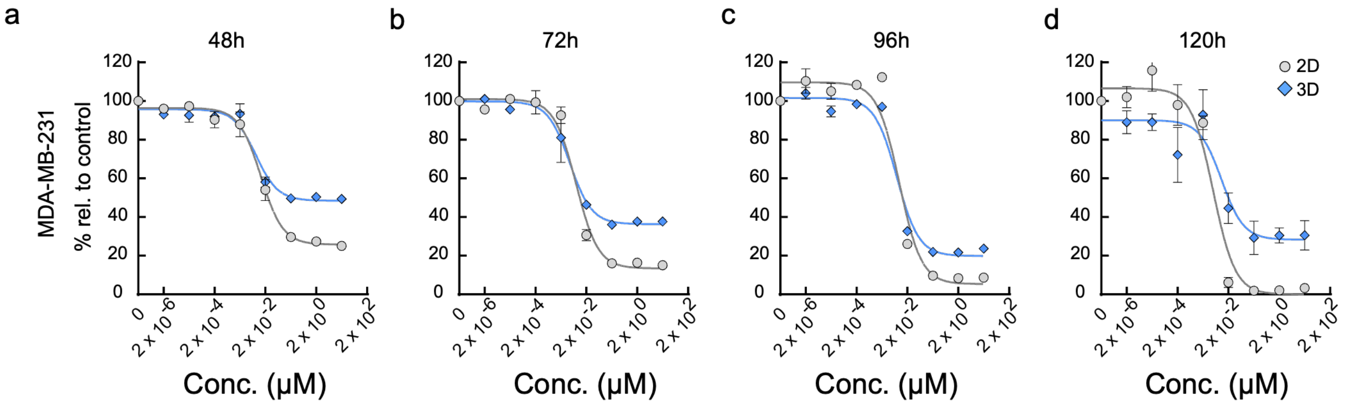


**Figure S4. Time-dependent taxol response of MDA-MB-231 cultured in 2D and 3D microenvironments.** Dose–response curves for MDA-MB-231 cultured in 2D or 3D and treated with increasing concentrations of taxol for (**a**) 48, (**b**) 72, (**c**) 96, and (**d**) 120 h. Responses are expressed as a percentage relative to the corresponding untreated control. Data points represent the mean ± s.e.m. from three independent biological replicates (n = 3). Dose–response curves were fitted using a four-parameter logistic regression model in GraphPad Prism v10.

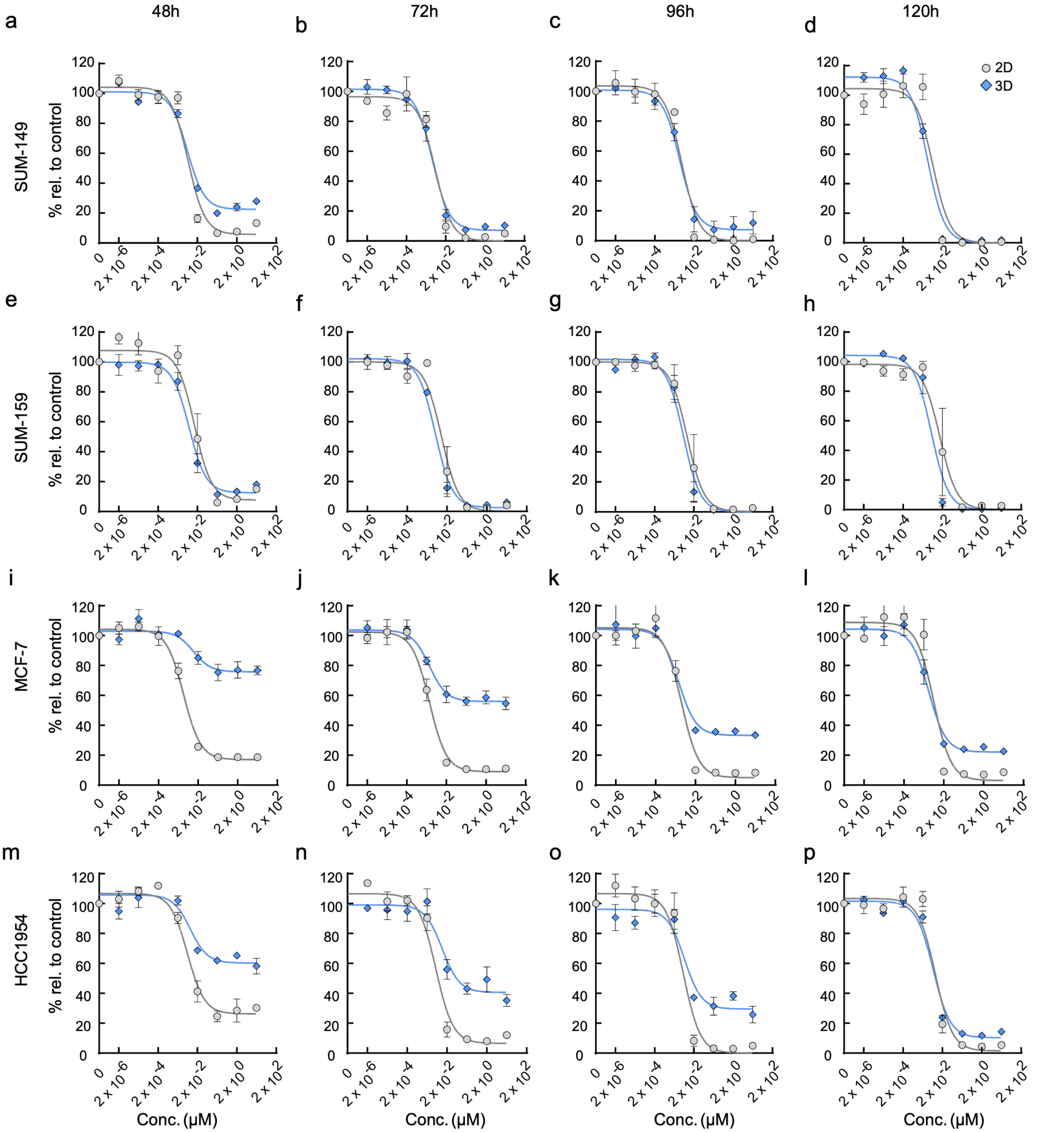


**Figure S5. Time-dependent taxol responses of breast cancer cell lines cultured under 2D and 3D conditions.** Dose–response curves for SUM-149 (**a–d**), SUM-159 (**e–h**), MCF-7 (**i–l**), and HCC1954 (**m–p**) cell lines cultured in 2D or 3D and treated with increasing concentrations of taxol for 48, 72, 96, or 120 h. Responses are expressed as a percentage relative to the corresponding untreated control. Data points represent the mean ± s.e.m. from three independent biological replicates (n = 3). Dose–response curves were fitted using a four-parameter logistic regression model in GraphPad Prism v10.

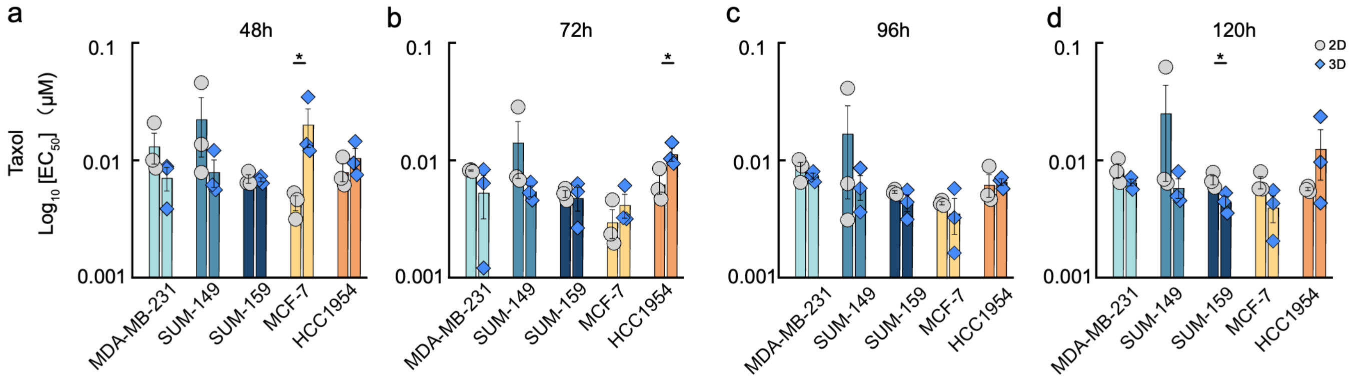


**Figure S6. Comparison of taxol $EC_{50}$ values among breast cancer cell lines cultured under 2D and 3D conditions.** $EC_{50}$ values for MDA-MB-231, SUM-149, SUM-159, MCF-7, and HCC1954 cell lines following taxol treatment for (**a**) 48, (**b**) 72, (**c**) 96, and (**d**) 120 h. $EC_{50}$ values were derived from four-parameter logistic dose–response curves fitted in GraphPad Prism v10 and are displayed on a logarithmic scale. Bars show the mean ± s.e.m., with individual data points shown for three independent experiments (n = 3). Statistical significance was assessed using two-tailed Student's *t*-tests.. (* $p \leq 0.05$).

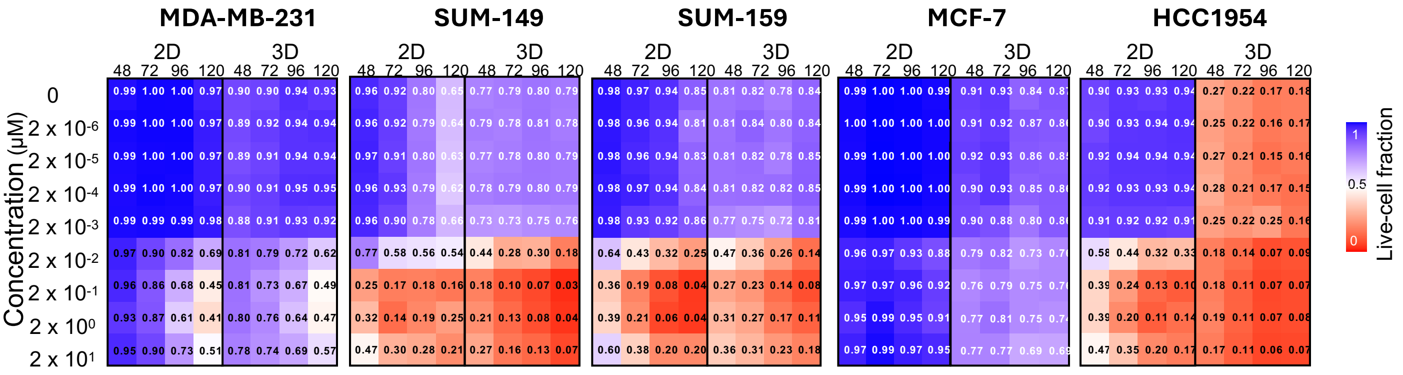

**Figure S7. Live-cell fraction across drug concentrations and taxol treatment durations in breast cancer cells cultured under 2D and 3D conditions.** Heatmaps show the live cell fraction, calculated as the number of live cells divided by the total number of cells (live + dead), following taxol treatment with increasing concentrations for 48, 72, 96, and 120 h in MDA-MB-231, SUM-149, SUM-159, MCF-7, and HCC1954 cell lines cultured in 2D or 3D. Values range from 0 (all cells dead) to 1 (all cells viable). Unlike conventional viability assays, which normalize cell viability to untreated controls and therefore reflect changes in total cell number relative to drug-free growth, this metric quantifies the proportion of surviving cells within each treatment condition independently of the untreated control, enabling direct assessment of treatment-induced net cell loss while minimizing the influence of differences in proliferation between culture conditions.

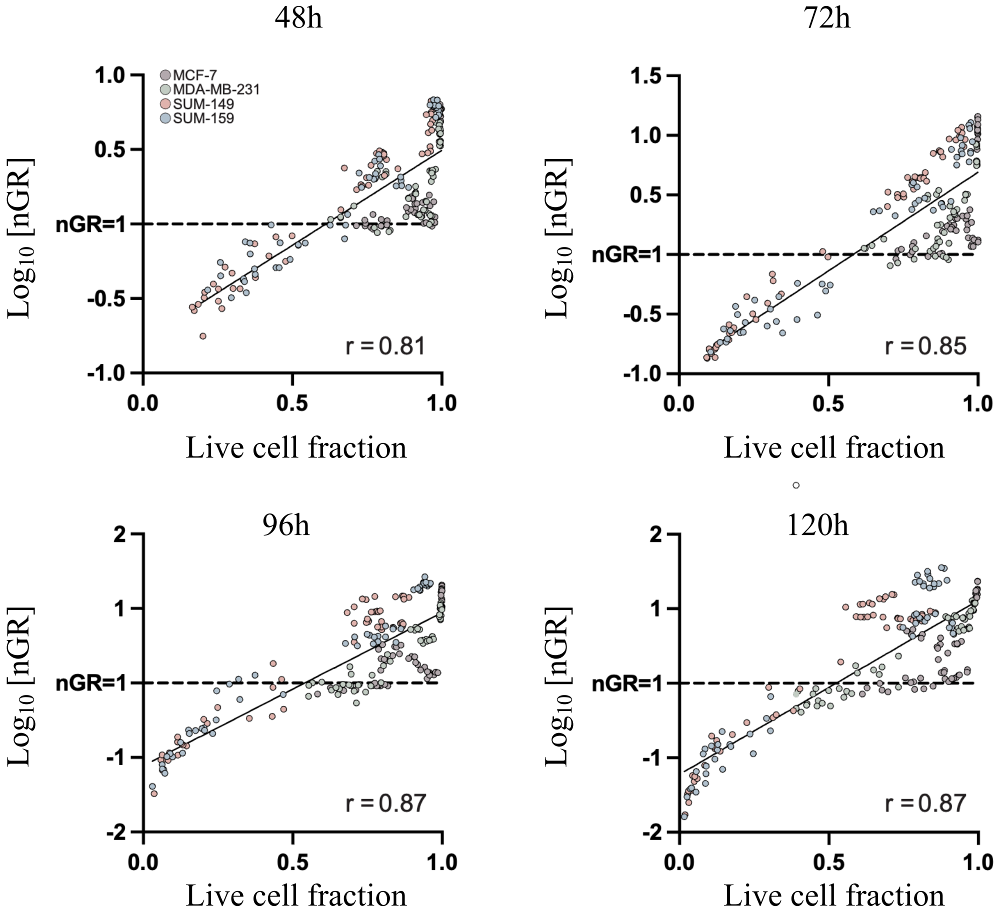


**Figure S8 Relationship between normalized growth ratio (nGR) and lice cell fraction (viability) across treatment durations.** Scatter plots showing the relationship between the live-cell fraction and log-transformed normalized growth ratio (nGR) at 48, 72, 96, and 120 h across MCF-7, MDA-MB-231, SUM-149, and SUM-159 cell lines under the tested treatment conditions. Each point represents an individual treatment condition, with colours indicating different cell lines. Solid lines indicate linear regression fits, and *r* denotes the Pearson correlation coefficient. The horizontal dashed line indicates nGR = 1 (log[nGR] = 0), corresponding to the boundary between net population growth (nGR > 1) and net population reduction (nGR < 1).

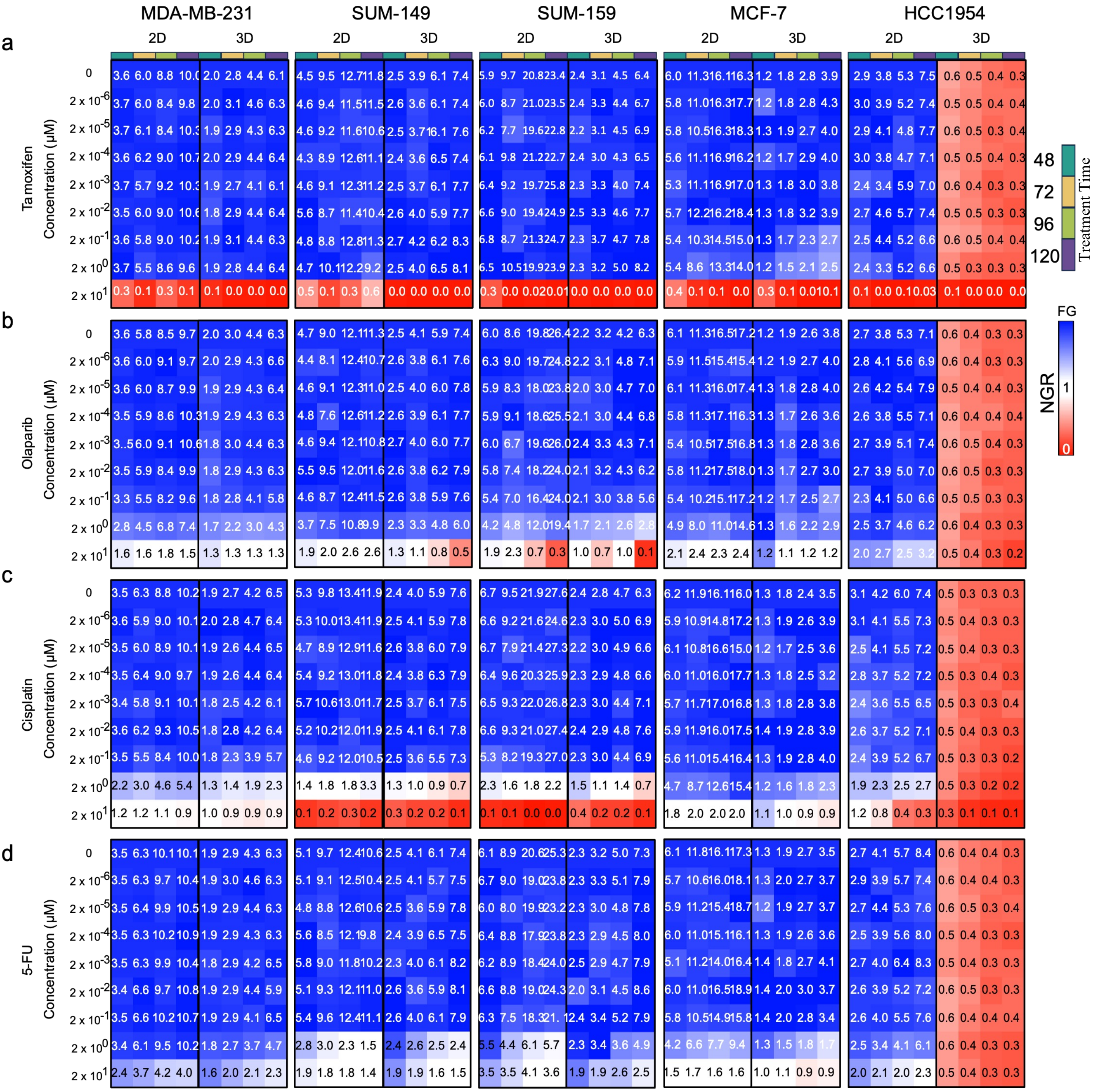


**Figure S9. Drug concentration- and time-dependent growth responses of breast cancer cell lines cultured in 2D and 3D.** Heatmaps showing normalized growth ratio (nGR) values across drug concentrations and time points (48, 72, 96, and 120 h) for five breast cancer cell lines, MCF-7, MDA-MB-231, SUM-149, SUM-159, and HCC1954, cultured in 2D and 3D and treated with (**a**) Tamoxifen, (**b**) Olaparib, (**c**) Cisplatin, and (**d**) 5-fluorouracil (5-FU). Colors indicate growth states: blue, uninhibited growth; white, partial growth inhibition; red, net cell loss. Values represent the mean of three independent experiments (n = 3).

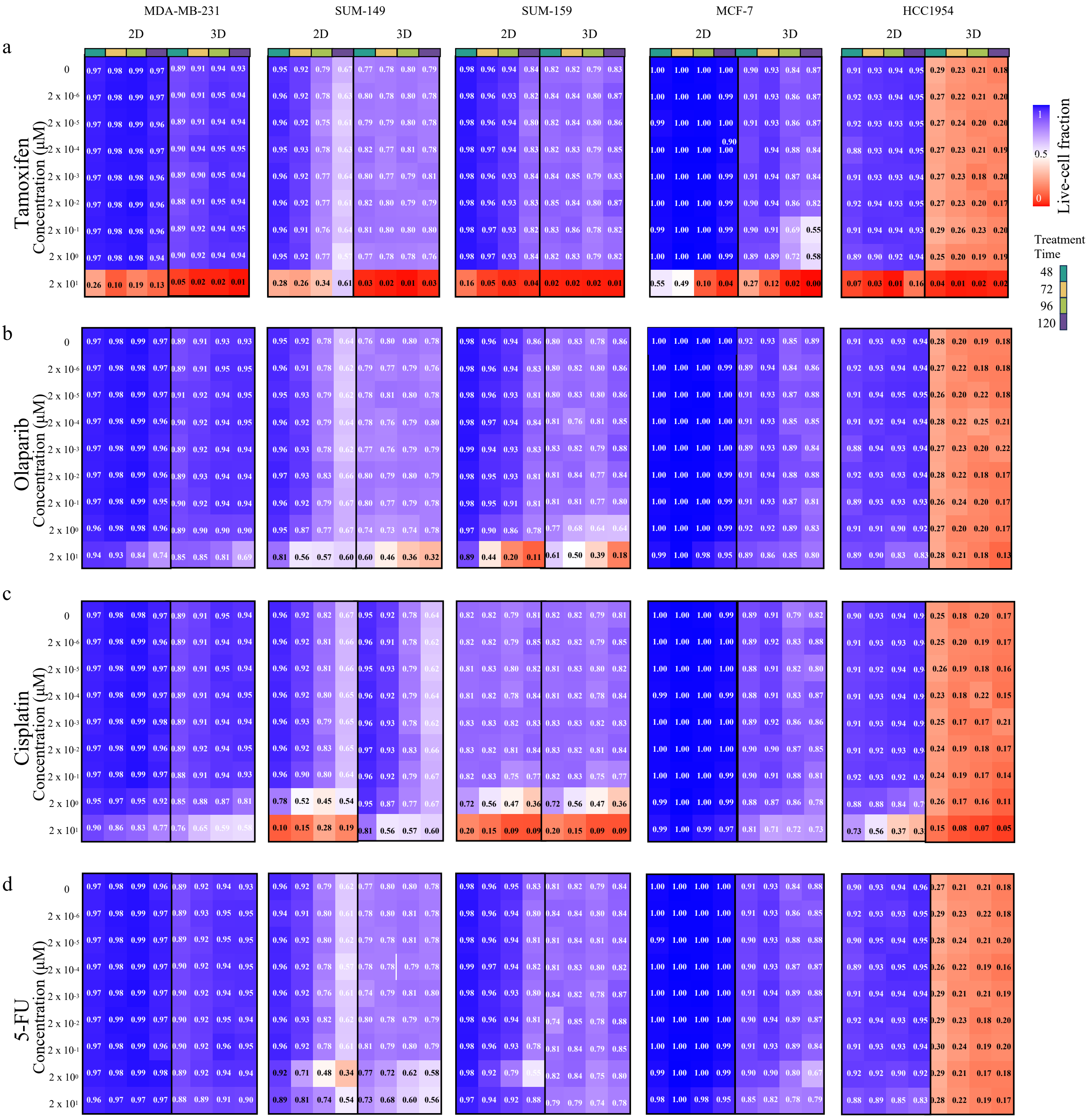

**Figure S10. Live-cell fraction across treatment conditions in 2D and 3D culture models**. Heatmaps showing the percentage of live cells across drug concentrations and time points (48, 72, 96, and 120 h) for five breast cancer cell lines, MCF-7, MDA-MB-231, SUM-149, SUM-159, and HCC1954, cultured in 2D and 3D and treated with (**a**) Tamoxifen, (**b**) Olaparib, (**c**) Cisplatin, and (**d**) 5-FU. The percentage of live cells was calculated as live cells/(live cells + dead cells). Colors indicate the percentage of live cells, with red representing lower and blue representing higher cell viability. Values represent the mean of three independent experiments (n = 3).

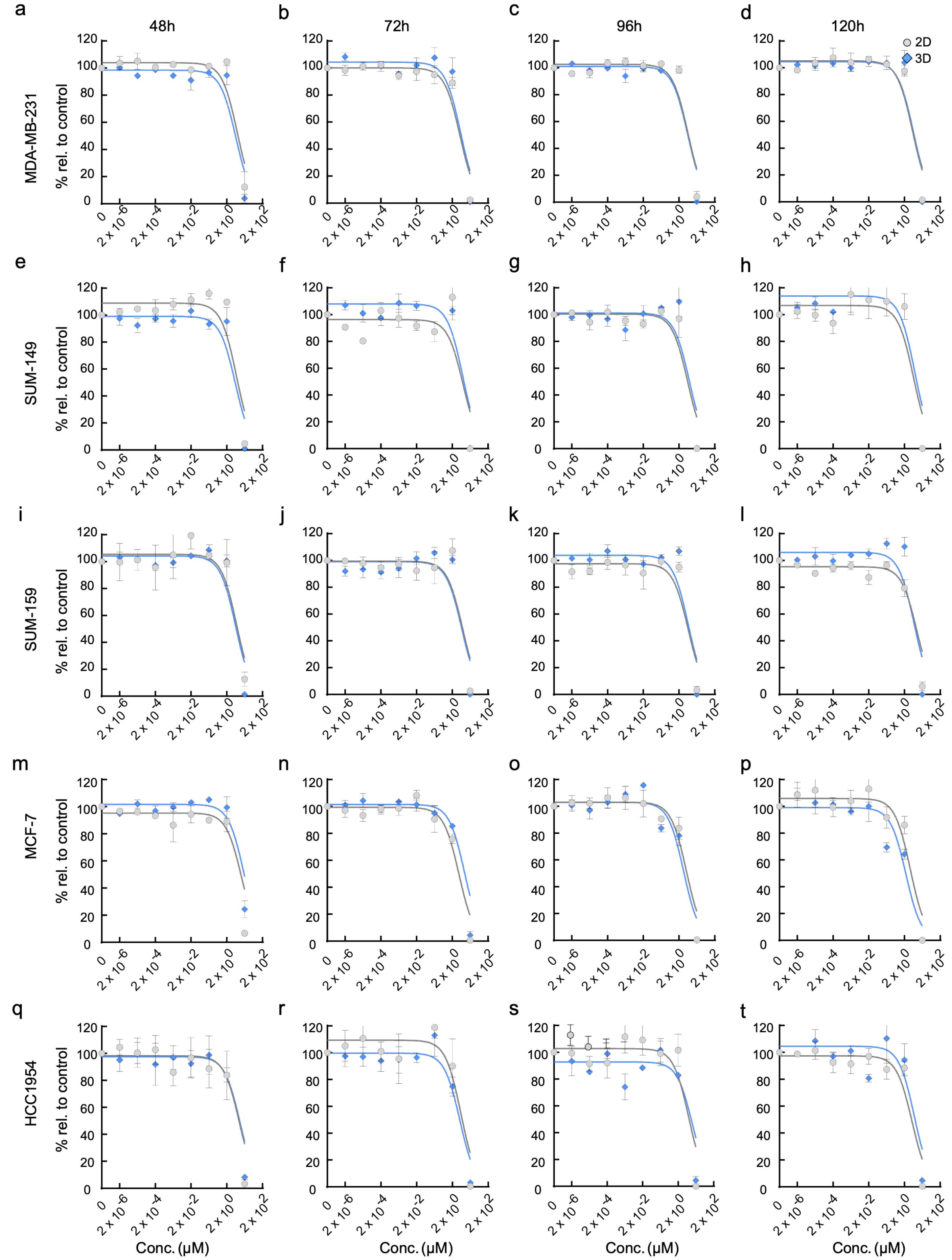


**Figure S11. Time-dependent tamoxifen responses of breast cancer cell liness cultured under 2D and 3D conditions.** Dose–response curves for MDA-MB-231 (**a–d**), SUM-149 (**e–h**), SUM-159 (**i–l**), MCF-7 (**m–p**), and HCC1954 (**q-t**) cell lines cultured in 2D or 3D and treated with increasing concentrations of tamoxifen for 48, 72, 96, or 120 h. Responses are expressed as a percentage relative to the corresponding untreated control. Data points represent the mean ± s.e.m. from three independent biological replicates (n = 3). Dose–response curves were fitted using a four-parameter logistic regression model in GraphPad Prism v10.

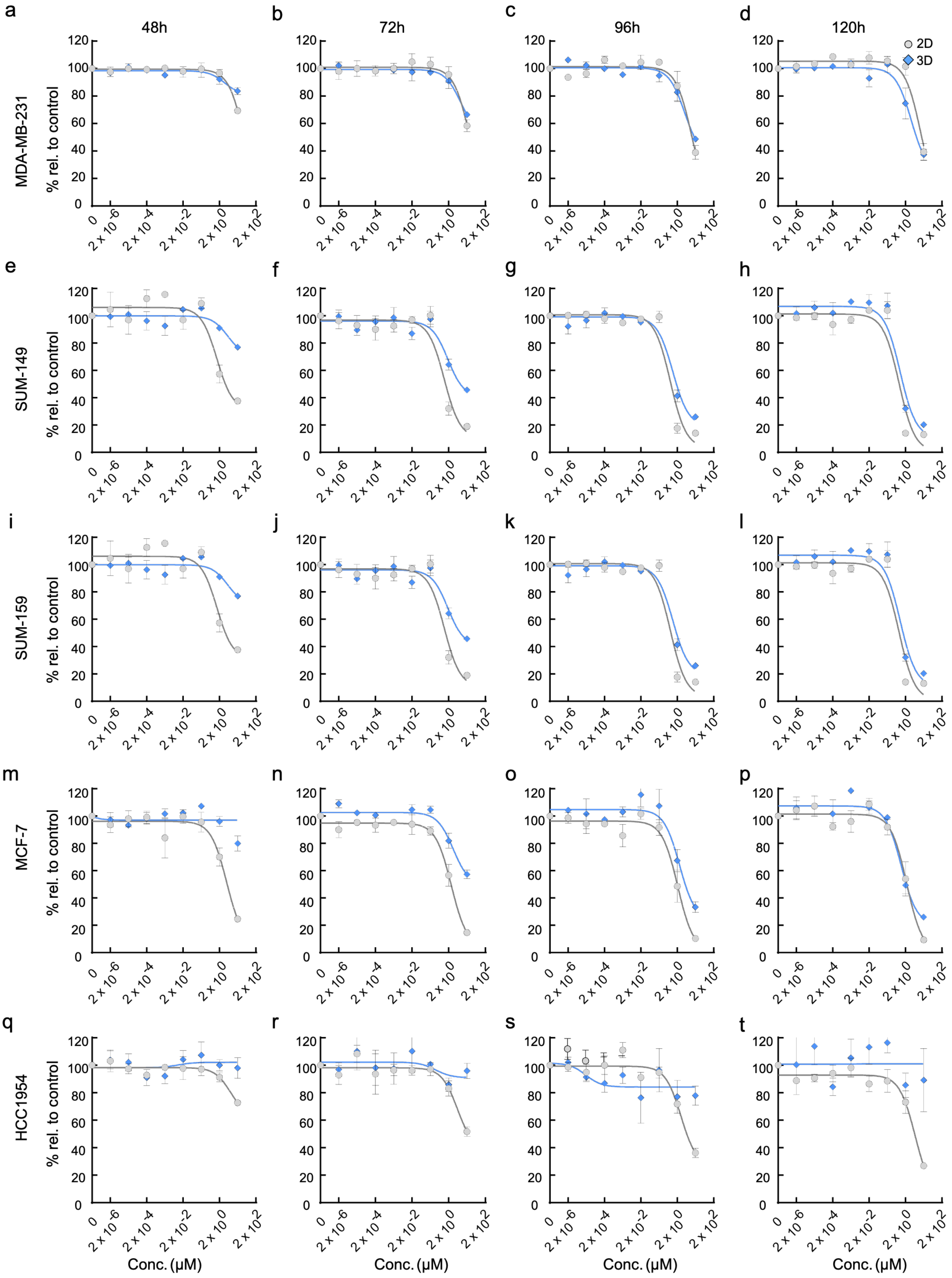


**Figure S12. Time-dependent olaparib responses of breast cancer cell lines cultured under 2D and 3D conditions.** Dose–response curves for MDA-MB-231 (**a–d**), SUM-149 (**e–h**), SUM-159 (**i–l**), MCF-7 (**m–p**), and HCC1954 (**q-t**) cell lines cultured in 2D or 3D and treated with increasing concentrations of olaparib for 48, 72, 96, or 120 h. Responses are expressed as a percentage relative to the corresponding untreated control. Data points represent the mean ± s.e.m. from three independent biological replicates (n = 3). Dose–response curves were fitted using a four-parameter logistic regression model in GraphPad Prism v10.

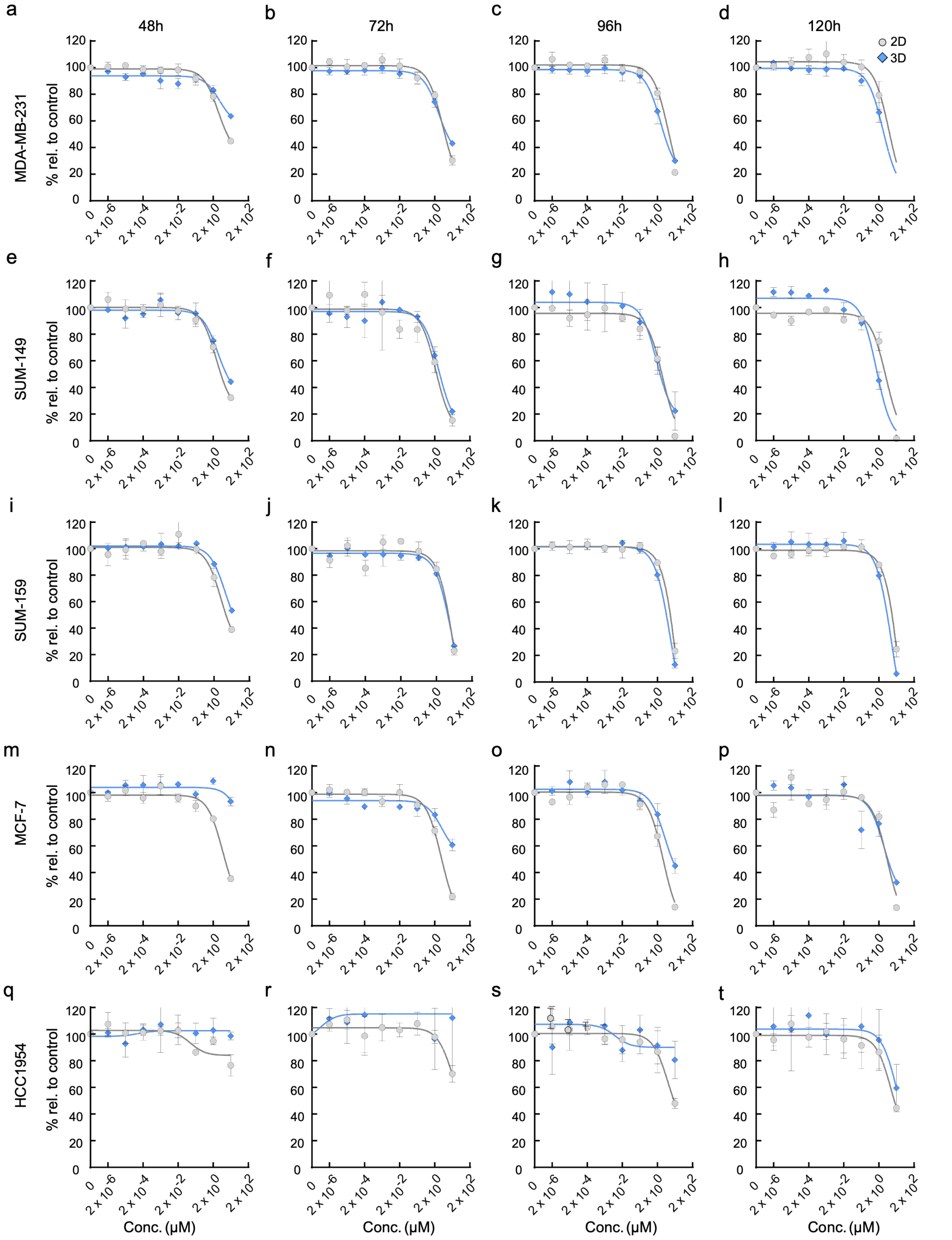


**Figure S13. Time-dependent cisplatin responses of breast cancer cell lines cultured under 2D and 3D conditions.** Dose–response curves for MDA-MB-231 (**a–d**), SUM-149 (**e–h**), SUM-159 (**i–l**), MCF-7 (**m–p**), and HCC1954 (**q–t**) cell lines cultured in 2D or 3D and treated with increasing concentrations of cisplatin for 48, 72, 96, or 120 h. Cell growth is expressed as a percentage relative to the corresponding untreated control. Grey circles represent 2D cultures, and blue diamonds represent 3D cultures. Data points represent the mean ± s.e.m. from three independent biological replicates (n = 3). Dose–response curves were fitted using a four-parameter logistic regression model in GraphPad Prism.

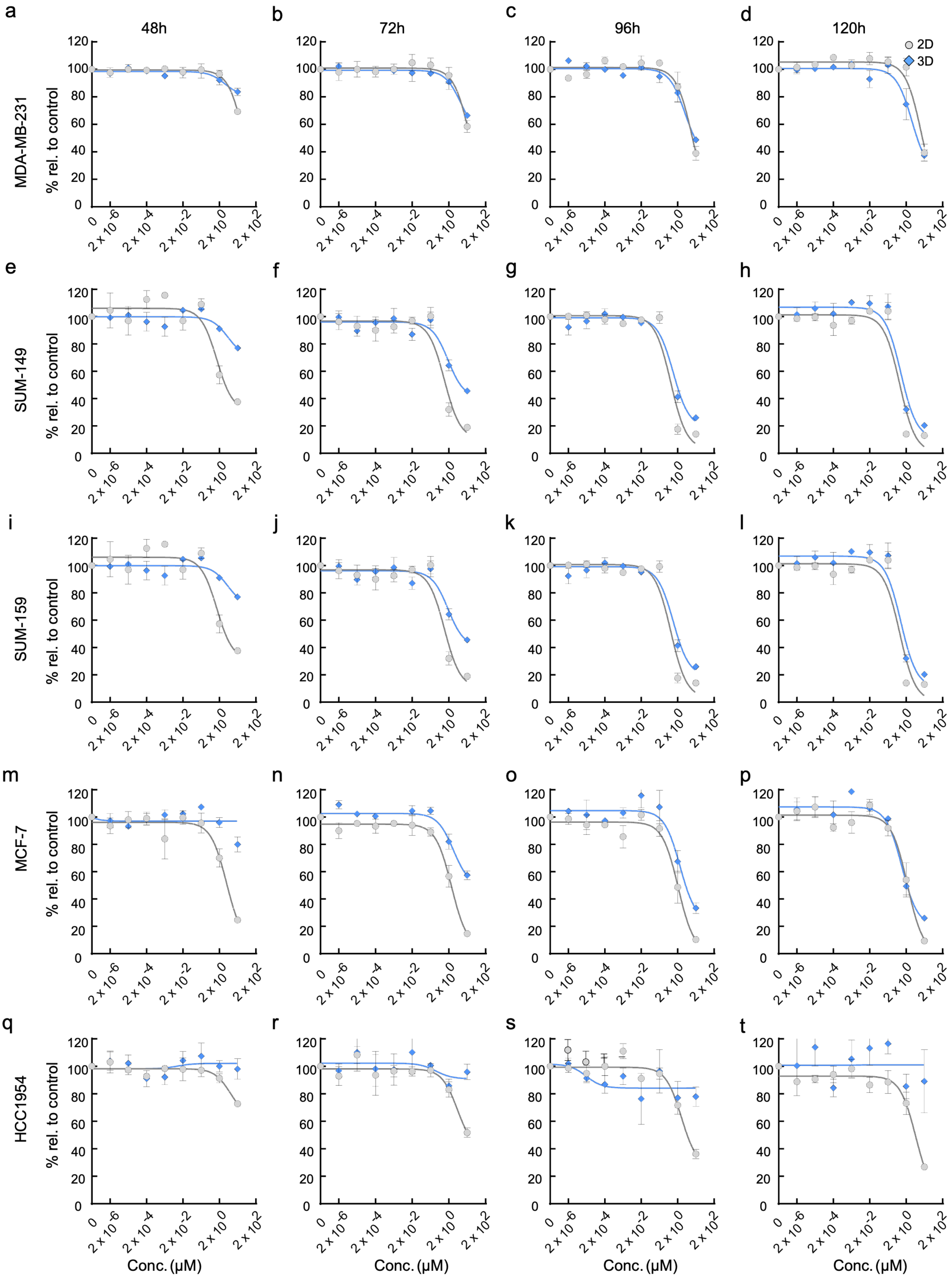


**Figure S14. Time-dependent 5-FU responses of breast cancer cell lines cultured under 2D and 3D conditions.** Dose–response curves for MDA-MB-231 (**a–d**), SUM-149 (**e–h**), SUM-159 (**i–l**), MCF-7 (**m–p**), and HCC1954 (**q–t**) cell lines cultured in 2D or 3D and treated with increasing concentrations of 5-FU for 48, 72, 96, or 120 h. Cell growth is expressed as a percentage relative to the corresponding untreated control. Grey circles represent 2D cultures, and blue diamonds represent 3D cultures. Data points represent the mean ± s.e.m. from three independent biological replicates (n = 3). Dose–response curves were fitted using a four-parameter logistic regression model in GraphPad Prism.

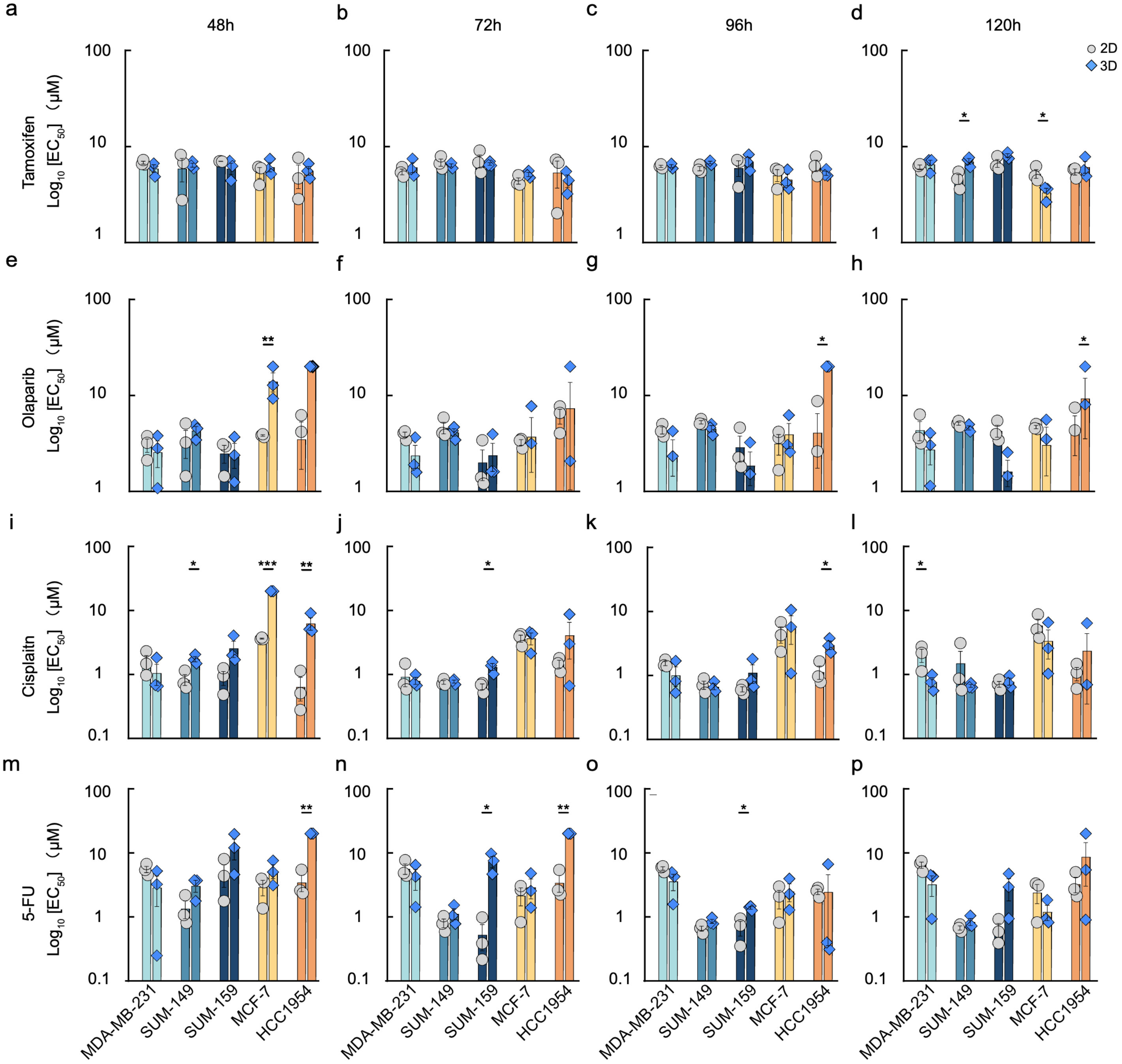


**Figure S15. Comparison of drug EC₅₀ values between breast cancer cell lines cultured under 2D and 3D conditions.** $EC_{50}$ values for MDA-MB-231, SUM-149, SUM-159, MCF-7, and HCC1954 cell lines following treatment with tamoxifen (**a–d**), olaparib (**e–h**), cisplatin (**i–l**), or 5-FU (**m–p**) for 48, 72, 96, or 120 h. $EC_{50}$ values were derived from four-parameter logistic dose–response curves fitted in GraphPad Prism and are presented on a logarithmic scale. Bars show the log (mean) ± s.e.m., with individual data points shown for three independent experiments (n = 3). Statistical comparisons were performed between the corresponding 2D and 3D conditions. *** $P < 0.001$; ** $P < 0.01$; * $P < 0.05$;

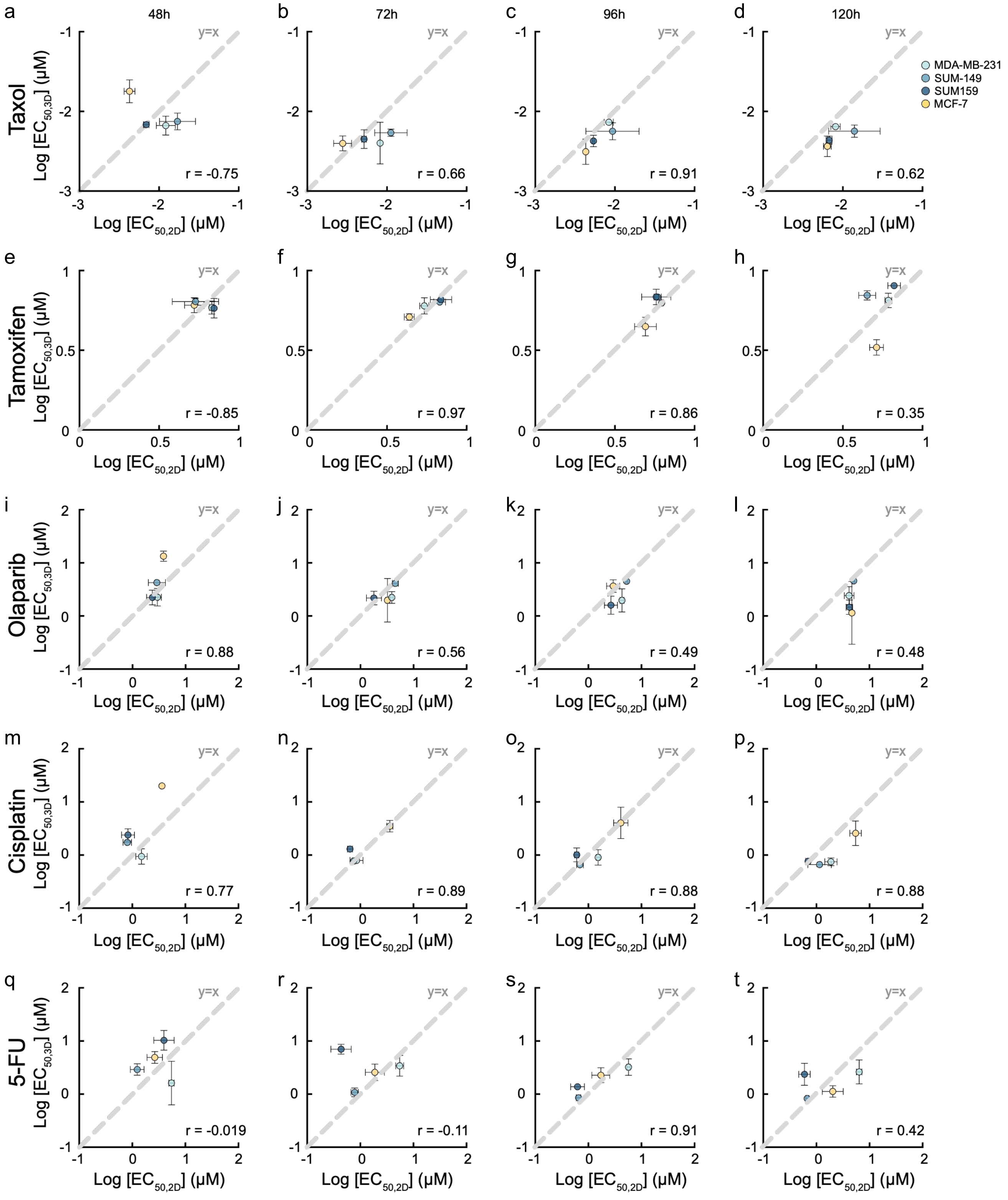


**Figure S16. Correlation of $EC_{50}$ between 2D and 3D breast cancer cultures across multiple treatment durations.** Correlation plots comparing log-transformed $EC_{50}$ values obtained from 2D and 3D cultures following treatment with taxol (**a–d**), tamoxifen (**e–h**), olaparib (**i–l**), cisplatin (**m–p**), and 5-FU (**q–t**) for 48, 72, 96, or 120 h. The dashed diagonal line represents equal concentration in 2D and 3D cultures (y = x). Each point represents the mean $EC_{50}$ value from three independent biological replicates, with error bars indicating s.e.m. Pearson's correlation coefficient (r) is shown for each treatment time**.**

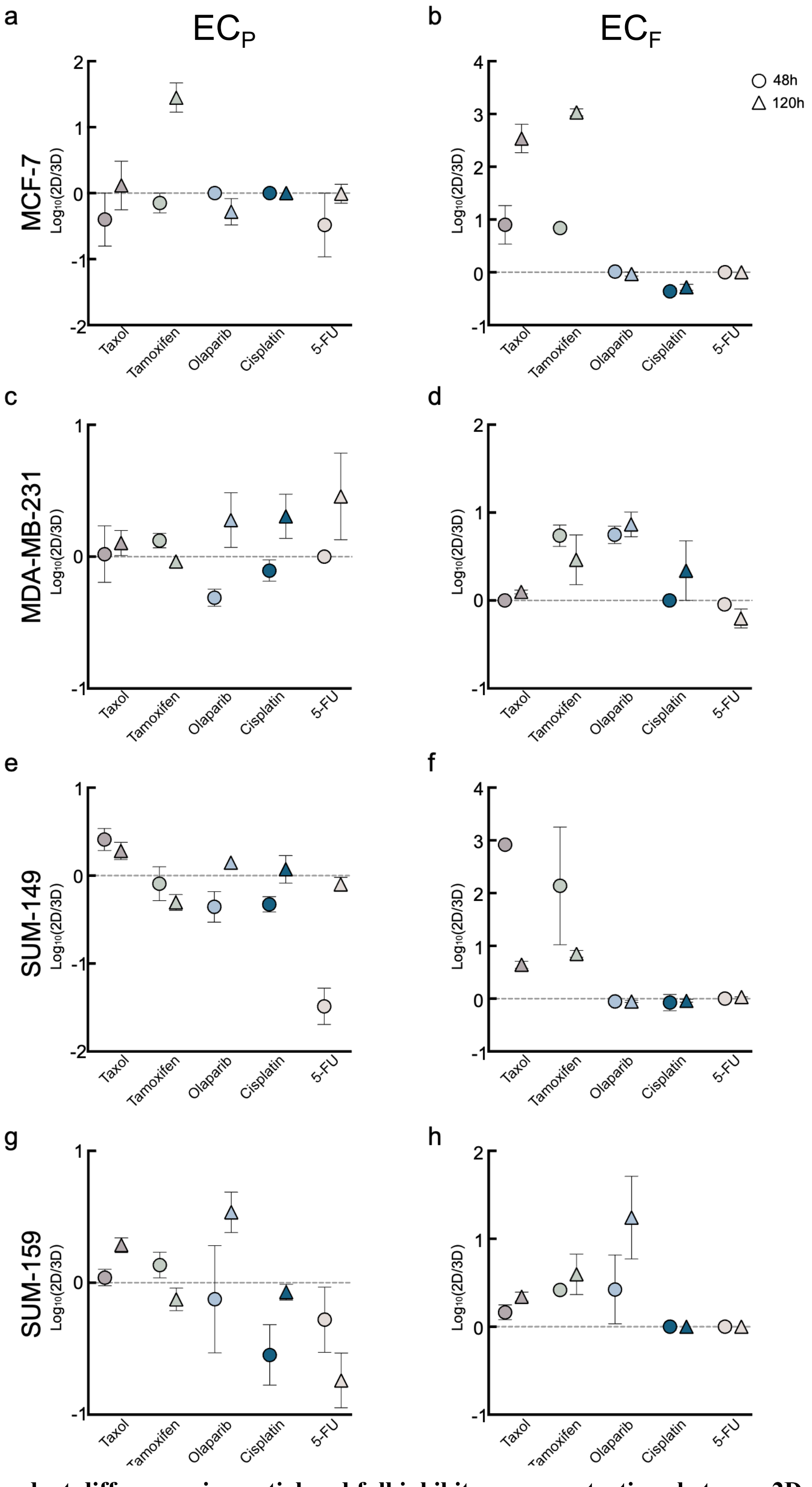


**Figure S17. Time-dependent differences in partial and full inhibitory concentrations between 2D and 3D cultures.** Log-transformed 2D-to-3D response ratios were calculated for partial inhibitory concentration ($EC_P$) and full inhibitory concentration ($EC_F$) values across treatment time. Representative plots are shown for four breast cancer cell lines treated with taxol, tamoxifen, olaparib, cisplatin, or 5-FU. $EC_P$ values are shown for MCF-7 (**a**), MDA-MB-231 (**c**), SUM-149 (**e**), and SUM-159 (**g**). $EC_F$ values are shown for MCF-7 (**b**), MDA-MB-231 (**d**), SUM-149 (**f**), and SUM-159 (**h**). Circles represent 48-h and triangles represent 120-h treatment. Data points represent the mean ± s.e.m. from three independent biological replicates (n = 3). The dashed line at 0 indicates equivalent responses between 2D and 3D cultures.

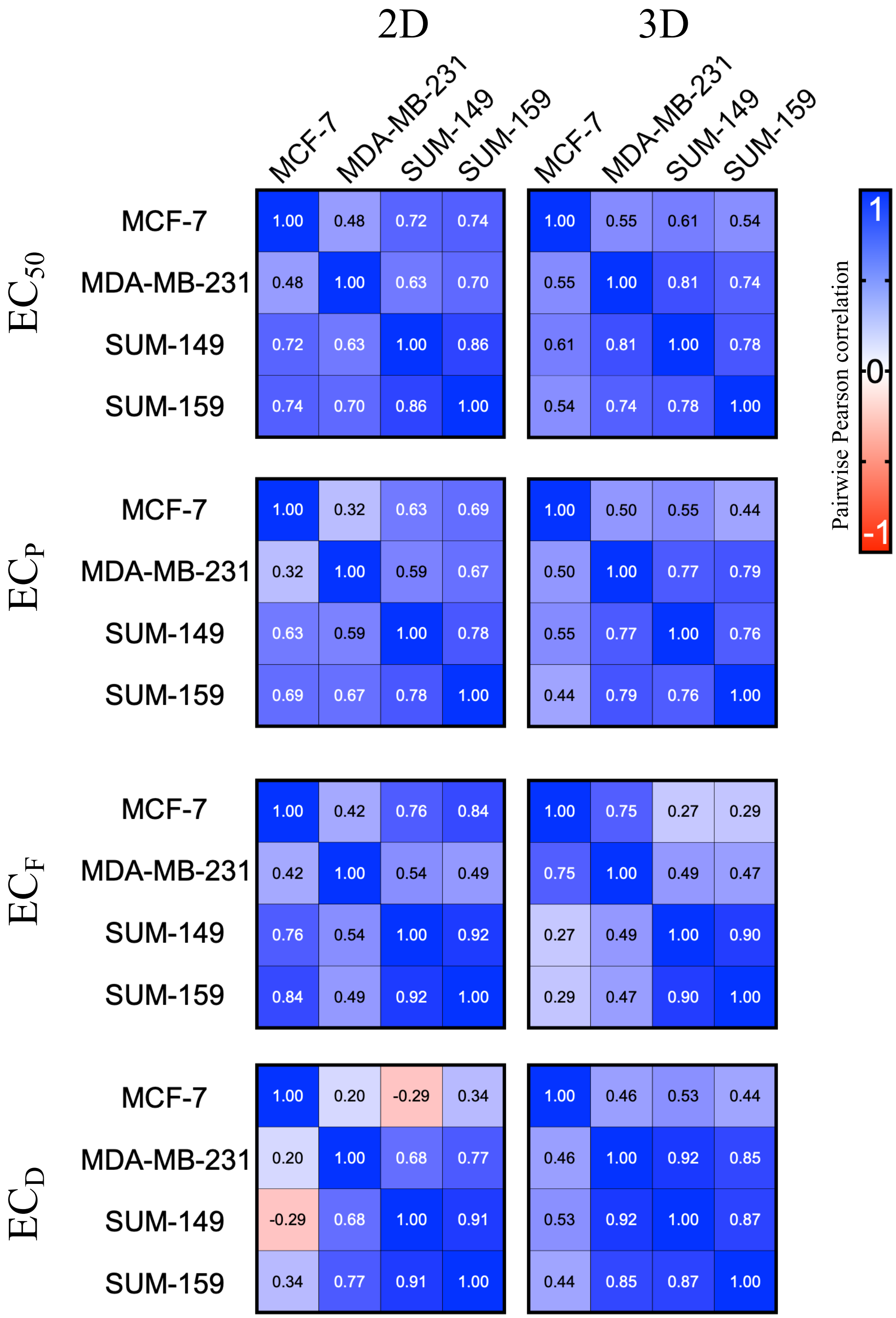


**Figure S18. Correlation of drug response metrics among breast cancer cell lines cultured under 2D and 3D conditions.** Pairwise Pearson correlation coefficients of drug response metrics across MCF-7, MDA-MB-231, SUM-149, and SUM-159 cell lines cultured in 2D (left) or 3D (right). Correlation matrices are shown for $EC_{50}$, $EC_P$, $EC_F$, and $EC_D$, calculated from responses to five anticancer drugs measured over four treatment durations (48, 72, 96, and 120 h). Numbers within each cell indicate the Pearson correlation coefficient (r).
